# E2VD: a unified evolution-driven framework for virus variation drivers prediction

**DOI:** 10.1101/2023.11.27.568815

**Authors:** Zhiwei Nie, Xudong Liu, Jie Chen, Zhennan Wang, Yutian Liu, Haorui Si, Tianyi Dong, Fan Xu, Guoli Song, Yu Wang, Peng Zhou, Wen Gao, Yonghong Tian

**Affiliations:** School of Electronic and Computer Engineering, Peking University, China; Peng Cheng Laboratory, Shenzhen, China; School of Computer Science, Peking University, China; CAS Key Laboratory of Special Pathogens, Wuhan Institute of Virology, Chinese Academy of Sciences, Wuhan, China; University of Chinese Academy of Sciences, Beijing, China; Guangzhou Laboratory, Guangzhou, China

## Abstract

The increasing frequency of emerging viral infections necessitates a rapid human response, highlighting the cost-effectiveness of computational methods. However, existing computational approaches are limited by their input forms or incomplete functionalities, preventing a unified prediction of diverse viral variation drivers and hindering in-depth applications. To address this issue, we propose a unified evolution-driven framework for predicting virus variation drivers, named E2VD, which is guided by virus evolutionary traits priors. With evolution-inspired design, E2VD comprehensively and significantly outperforms state-of-the-art methods across various virus variation drivers prediction tasks. Moreover, E2VD effectively captures the fundamental patterns of virus evolution. It not only distinguishes different types of mutations but also accurately identifies rare beneficial mutations that are critical for virus to survival, while maintains generalization capabilities on different viral lineages. Importantly, with predicted biological drivers, E2VD perceives virus evolutionary trends, in which potential high-risk mutation sites are accurately recommended. Overall, E2VD represents a unified, structure-free, and interpretable approach for analyzing and predicting viral evolutionary fitness, providing an ideal alternative to costly wet-lab measurements to accelerate responses to emerging viral infections.

## 1 Introduction

Since 2000, numerous emerging viral infections [1, 2], especially COVID-19 [3], have seriously threatened the global public health system. Severe acute respiratory syndrome coronavirus 2 (SARS-CoV-2), the causative agent of COVID-19, infiltrates cells by targeting the human angiotensin-converting enzyme 2 (ACE2) as the host receptor [4–6]. To combat the ongoing pandemic, various vaccines and drugs have been developed to mitigate the damage caused by SARS-CoV-2 [7, 8]. However, a concerning aspect is that SARS-CoV-2 continues to evolve, with mutations accumulating and recombination processes contributing to increased diversity [9, 10]. The sub-lineages of Omicron, in particular, remain prevalent and continue to cause breakthrough infections [11].

The outbreak of SARS-CoV-2 is only a microcosm of plenty emerging viral infections, reminding us to always pay attention to the possible emergence of unknown pathogens. Furthermore, the rising occurrence of emerging viral infections necessitates faster reactions from the human. Uncovering biological drivers to analyze and predict evolutionary trends is an effective approach to increase response speed. Correspondingly, some wet-lab techniques have been developed to measure the mutation-induced properties of viruses [12–15], which are then adopted to analyze the evolutionary trajectories and mutation hot-spots. However, wet-lab measurements are often used for post-hoc analysis due to their huge cost of money and time, preventing them from being carried out as an active-exploration approach. Therefore, there is an urgent requirement for efficient and low-cost computational methods to explore evolutionary trajectory of virus.

The computational methods currently available for virus evolution analysis, especially deep-learning models, can be categorized into two types, i.e. structure-dependent methods that requires structural information and sequence-only methods based only on sequences. Structure-dependent methods [16–19] require the tertiary structure of proteins as input, such as antibodies, the ACE2, or various protein complexes. In contrast, sequence-only methods [20–26] adopt protein sequences as the only input information. The above methods can also be divided into three types in terms of their functions, including specific property prediction, evolutionary trajectory reproduction, and evolutionary trend prediction. Methods for specific property prediction [18, 19, 21, 22, 24–26] aim to predict mutation-induced viral protein property changes, such as binding affinity, expression, and antibody escape. The evolutionary trajectory reproduction methods [20] can reproduce the evolutionary route of the observed variants. The evolutionary trend prediction methods [16, 17, 23] are capable of providing predictive analysis of potential evolutionary trends.

However, current methods have significant limitations. For structure-dependent methods [16–19], they are severely limited by the rarity or even absence of high-quality protein structures within specific families. Even though deep-learning-based protein structure prediction methods [27–30] have made significant progress, they struggle to deal with the viral protein sequences with a few mutations [31–36]. In contrast, one-dimensional protein sequences are easy to obtain, but the lack of structural and functional information is often the performance bottleneck of sequence-only methods [20–26]. In the aspect of functions, current specific property prediction methods [18, 19, 21, 22, 24–26] lack comprehensiveness and unification, and only develop one or two customized predictors for specific viral protein properties. Besides, current evolutionary trajectory reproduction methods [20] can only process observed variants and lacks predictive capability for future variants, which is essentially a type of post-hoc data modeling and analysis. While there exist several evolutionary trend prediction methods like those discussed in [16, 17, 23], they exhibit clear limitations, such as incomplete consideration of diverse mutational drivers, and dependence on structure input. Therefore, it is crucial to develop a unified, structure-free, and interpretable method for virus variation drivers prediction and apply it effectively to the prediction of evolutionary trends.

The prerequisite for achieving unification and interpretability is to design a deep-learning framework under the guidance of the fundamental principles of virus evolution. Mutation is the cornerstone of viral evolution [37], and RNA viruses have significantly higher mutation rates [32]. While a small proportion of mutations are beneficial and some are nearly neutral, the majority of mutations are harmful to viral fitness [31]. The ratio of harmful to beneficial mutations may vary across organisms and environments, but harmful mutations are thought to always outnumber beneficial mutations across the evolutionary scale [31, 32]. In other words, beneficial mutations are a rare subset of the viral protein’s evolutionary fitness [38]. Naturally, having numerous harmful mutations greatly decreases the probability of multiple mutations coexisting within a sequence. Generally, comparing with wild-type virus, only few sites in a viral protein sequence would mutate. As shown in Fig.1a, we can summarize the above priors as ‘rare beneficial mutations’ and ‘few-site mutations’, which would bring corresponding challenges for prediction. First, mutations at few sites are difficult to significantly affect the reconstruction of molecular interactions, which results in a very subtle effect on viral protein fitness, making it difficult for neural networks to capture. Second, the scarcity of beneficial mutations causes extreme imbalance in sample categories from a data perspective, which poses a huge challenge to training deep learning models that accurately mine rare beneficial mutations.

**Fig. 1.**
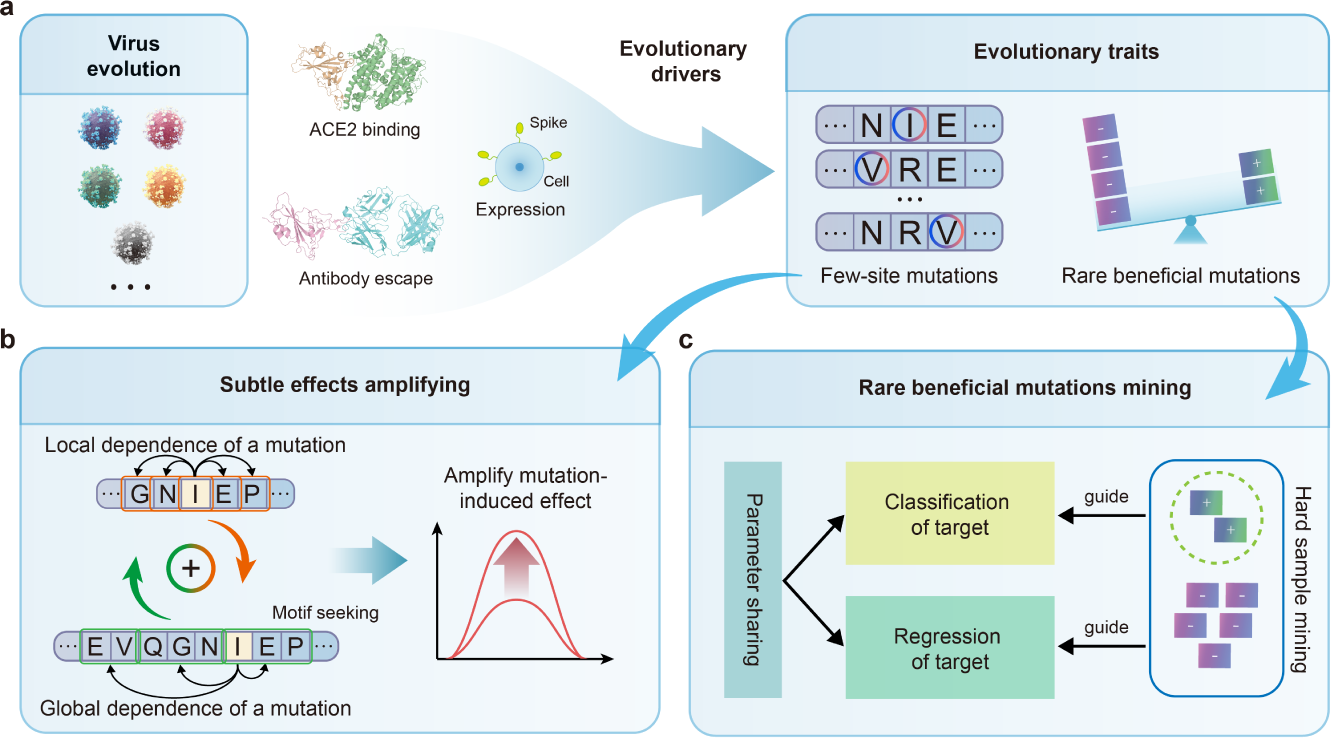
The motivation and methodology of E2VD. **a**, The evolutionary traits of viruses under diverse evolutionary biological drivers, including ‘few-site mutations’ and ‘rare beneficial mutations’. **b**, Local-global dependence coupling module is proposed to amplify the subtle effects caused by mutations at few sites through integrating the local dependence of a mutation on nearby amino acids and the global dependence at the motif level over the entire sequence. **c**, Multi-task focal learning module is proposed to force the model to assign more focus on rare beneficial mutations to improve prediction performance through multi-task learning mode.

In order to solve the above challenges, we propose E2VD (**E**volution-driven **V**irus **V**ariation **D**rivers prediction), a unified deep-learning framework for predicting biological drivers for virus. We propose two innovative modules, local-global dependence coupling, and multi-task focal learning. The first innovative module, local-global dependence coupling, integrates the local dependence on nearby amino acids and the global dependence at the motif level over the entire sequence, thereby amplifying subtle effects caused by few-site mutations (Fig.1b). The second innovative module, multitask focal learning, aims to alleviate the challenge caused by the severe imbalance between beneficial and harmful mutations. As shown in Fig.1c, accurately predicting rare beneficial mutations can be considered as a problem of hard sample mining with beneficial mutations as hard samples. Prediction for a specific variation driver is converted into two tasks: classification and regression of a target viral protein property. The prediction performance is promoted by learning the above related tasks simultaneously under the framework of multi-task learning [39] with the supervision of a novel ‘multi-task focal loss’.

With the support of the above two innovative modules, E2VD predicts key virus variation drivers (including binding affinity, expression, and antibody escape) in a structure-free form, significantly outperforming the state-of-the-art methods across the board. Moreover, E2VD effectively captures the fundamental patterns of virus evolution. It not only clearly distinguishes different types of mutations that make up the evolutionary fitness landscape, but also accurately identifies rare beneficial mutations critical for virus survival. In addition, E2VD exhibits enhanced generalization ability than its competitors across different lineages, indicating its potential to cope with intricate evolutionary trends. Importantly, the binding affinity, expression and antibody escape predictors developed based on E2VD are flexibly combined to comprehensively examine the evolutionary fitness of variants, thereby accurately recommend potential high-risk mutation sites. Therefore, E2VD offers an alternative cost-effective *in silico* approach for analyzing and predicting viral evolutionary fitness, accelerating responses to emerging viral infections.

## 2 Results

### 2.1 The architecture of E2VD is driven by evolutionary traits

The overall architecture of E2VD is shown in Fig.2a, and its core is to solve the modeling problems caused by evolutionary traits. First, considering that protein language models (PLMs) have made impressive progress on functional and structural prediction tasks [29, 38, 40–43], we adopt pre-trained PLMs with frozen parameters to extract amino-acid-level embeddings of protein sequences, including mutated receptor-binding domains (RBDs) of SARS-CoV-2 and antibody variable-regions. Next, the above amino-acid-level embeddings are fed into the **L**ocal-**G**lobal dependence coupling (LG) module. In this module, Convolutional Neural Network (CNN) [44] is adopted to build local dependencies of mutations, while ContextPool-attention [45] is used to build global dependencies of mutations at the motif level. The purpose of using CNN here is to emphasize the impact of a mutation on their sequentially neighboring amino acids to establish the local interaction system. However, local dependence alone is not sufficient to simulate the complete intramolecular interaction system faced by mutations, as it may ignore the interactions caused by spatial proximity, which highlights the need to introduce global dependencies. To go a step further, considering that biologically meaningful protein fragments, i.e., motifs [46], carry functional information [47], we adopt ContextPool-attention with dynamic granularity to establish the global dependencies of mutations based on motifs, thereby complementing the local dependencies of mutations. Specifically, we set a learnable granularity to conduct phrase mining, with the identified phrases matching motifs found in protein sequences. The global dependencies established on this basis take into account motifs carrying specific spatial conformation or specific functional information, thereby introducing more explicit sequence patterns corresponding to functions. According to the above approach, the local dependencies and motif-level global dependencies are extracted simultaneously, and integrated to get refined protein-level embedding.

**Fig. 2.**
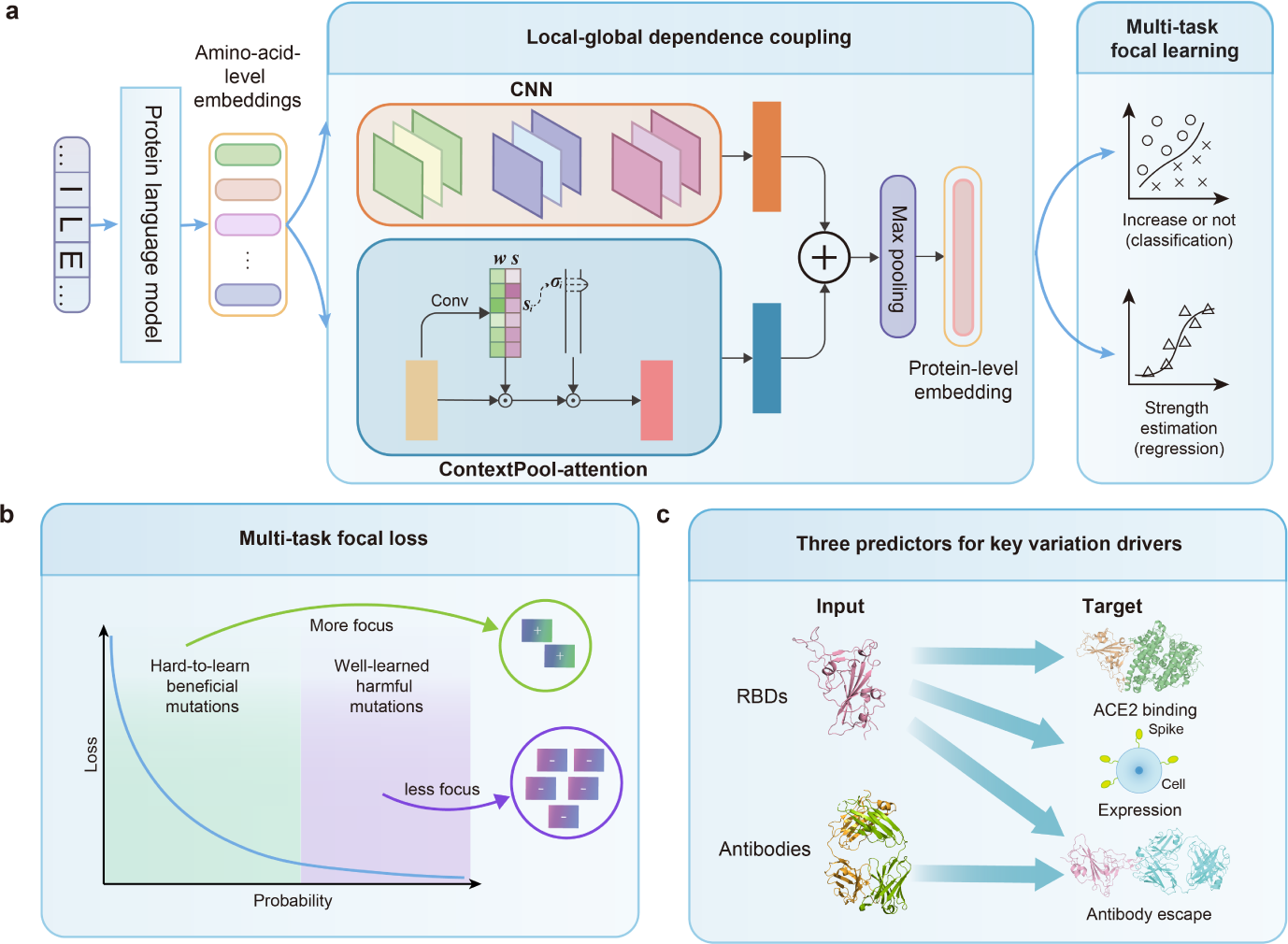
Model architecture and types of prediction tasks. **a**, The overall architecture of E2VD. A protein sequence is first extracted as amino-acid-level embeddings, and then converted into a protein-level embedding through the local-global dependence coupling module consisting of CNN and ContextPool-attention. Finally, the protein-level embedding flows into the multi-task focal learning module for multi-task learning of classification and regression. **b**, Multi-task focal loss is designed to help the model better learn hard-to-learn samples, i.e. rare beneficial mutations. Beneficial mutations are assigned more attention due to their rarity, while harmful mutations that are already well-learned in large numbers do not require much attention. **c**, Three types of prediction tasks for E2VD, including binding affinity, expression and antibody escape. The binding affinity and expression prediction tasks require the input of the mutated RBDs, while the antibody escape prediction task requires the input of both the mutated RBDs and the antibodies.

The refined protein-level embedding then flows into the **M**ulti-**T**ask focal learning (MT) module. In this module, the prediction of specific variation driver (i.e. viral protein property) is transformed into two closely related tasks, classification and regression, which is allowed for simultaneous training through multi-task learning. More importantly, hard-to-learn samples, i.e. rare beneficial mutations, require customized loss functions to ensure heightened focus by the neural network (Fig.2b). Focal loss for object detection [48] performs well in classification tasks with imbalanced categories, which inspired us to design customized focal loss for regression tasks. The focal loss for classification and designed focal loss for regression are combined into our proposed ‘multi-task focal loss’, which simultaneously incorporates task correlation and hard sample mining strategies.

The virus variation process is driven by a variety of factors, which requires E2VD to be unified to accommodate various prediction scenarios. In this work, we consider viral protein binding affinity, expression, and antibody escape as three key biological drivers [12, 49, 50] that significantly influence the intrapandemic evolution of SARS-CoV-2 (Fig.2c). The binding affinity and expression level are predicted based on mutated RBD sequences only, as the binding target (human ACE2) is fixed here, and the expression level only involves RBD sequences. Antibody escape prediction requires input for both RBD sequences and antibody sequences as it involves neutralization between variants and multiple antibodies.

### 2.2 Prediction performance of E2VD

There are currently a variety of off-the-shelf PLMs to choose from, such as SeqVec [43], ProtTrans series [40], ESM-2 series [29], and ESM-1v [41]. However, ESM-1v compared the impact of pre-training datasets with different sequence identities on zero-shot inference performance, highlighting the importance of pre-training data for PLMs, which inspires us to explore how to train specialized PLMs suitable for prediction of virus variation drivers. To improve adaptability for virus evolution, we customize the pretraining dataset selection and the truncation sequence length to create a proprietary PLM. First of all, the evolution of viruses involves the continuous accumulation of novel mutations [31, 32, 52], which requires the PLMs to have enough zero-shot generalization capability to maintain the quality of the extracted viral protein sequence embedding. We adopt the Uniref90 dataset which has been shown to yield optimal zero-shot inference performance by ESM-1v [41]. Furthermore, considering the relatively short sequence lengths of input RBDs (201) and antibody variable regions (136), we set the truncation sequence length for pre-training as 256. This adjustment aims to facilitate the identification of protein fragment patterns smaller than the specified length, thereby improving the model’s ability to capture essential protein features.

To evaluate the impact of customized pre-training configuration on the prediction performance, the widely recognized pre-trained PLMs are adopted to compare with our customized PLM on the binding affinity, expression, and antibody escape prediction task (Fig.3abc). To ensure a fair comparison, we only replace different pre-trained PLMs in the original complete E2VD architecture, and the remaining modules remain unchanged. E2VD with our customized PLM achieves the highest accuracy and Pearson’s Correlation Coefficient (PCC) with the smallest parameter size (340M), which proves the effectiveness of our customized pre-training configuration.

**Fig. 3.**
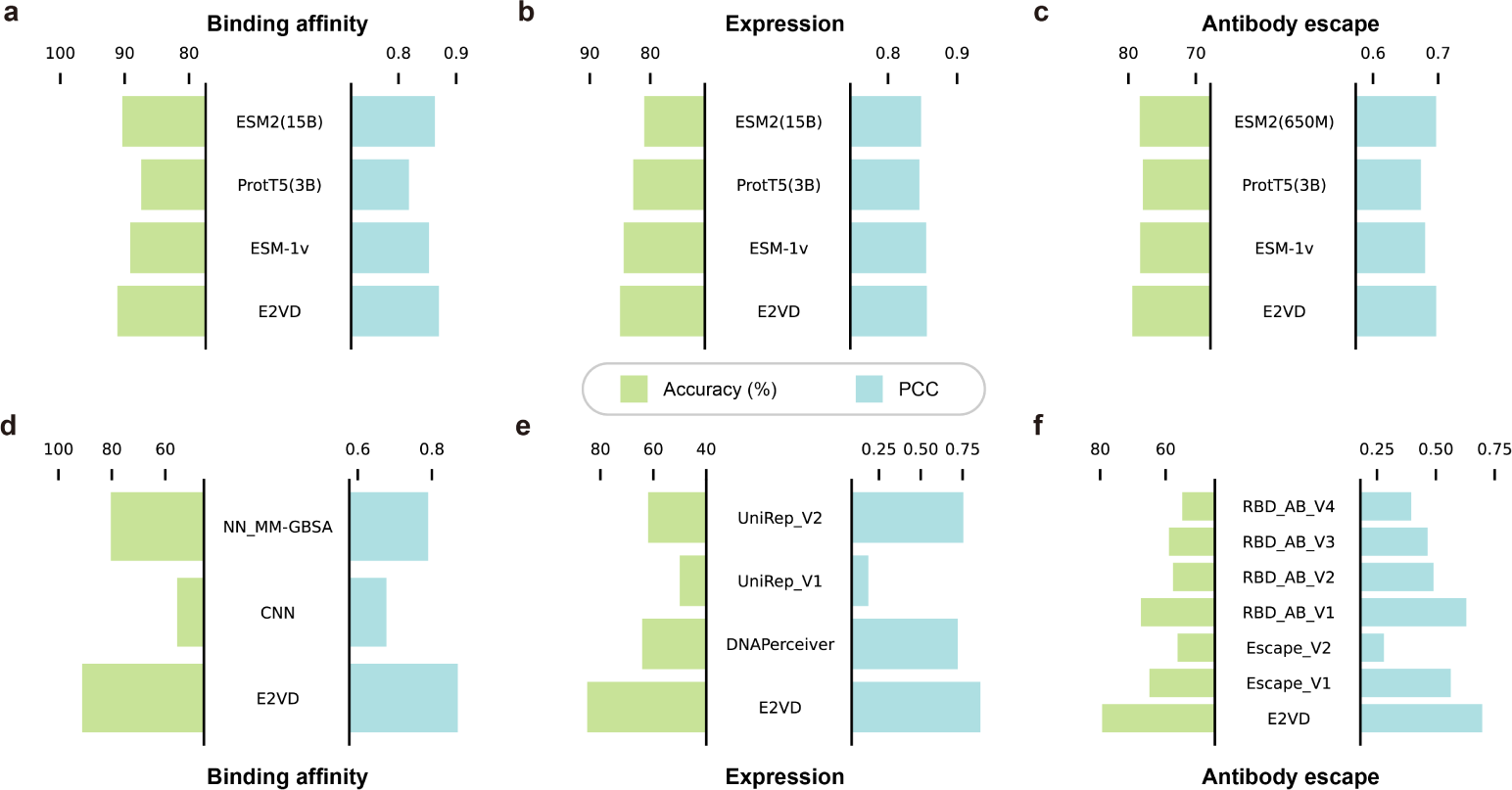
Prediction performance of E2VD. **abc**, Prediction performance of off-the-shelf PLMs and our customized one on three key variation drivers prediction tasks, including binding affinity (a), expression (b), and antibody escape (c). To ensure a fair comparison, we only replace different pretrained PLMs on the original complete architecture of E2VD, and the remaining modules remain unchanged. **def**, E2VD is benchmarked on binding affinity (d), expression (e), and antibody escape (f) prediction tasks, respectively. For expression prediction, two models of a previous study [25] trained with UniRep embeddings [51] for Random Forest (RF) and Artificial Neural Network (ANN) training are referred to as UniRep V1 and UniRep V2. For antibody escape prediction, two antibody escape prediction models of a previous study [22] based on pre-trained protein language models are referred to as Escape V1 and Escape V2. In addition, four models of the previous method RBD AB [26] with different masking ratios are referred to as RBD AB V1, RBD AB V2, RBD AB V3, and RBD AB V4, respectively.

As shown in Fig.3def, E2VD is benchmarked on binding affinity, expression, and antibody escape prediction tasks, respectively. First, for binding affinity prediction, the widely used CNN and a previous method NN MM-GBSA [19] are adopted for comparison. The accuracy of other methods in the classification task is improved by 10.7% by E2VD, along with a 8% improvement of the PCC in the regression task. Second, for expression prediction, three models from two previous studies [24, 25] are used for comparison. The accuracy of other methods in the classification task is improved by E2VD by 20.8%, and the PCC in the regression task is improved by 10.2%. Third, for antibody escape prediction, six models from two previous studies [22, 26] are used for comparison. The accuracy of other methods in the classification task is improved by more than 11.8% by E2VD, and the PCC in the regression task is improved by about 6.8%. Overall, E2VD comprehensively and significantly surpasses the state-of-the-art methods in the above three prediction tasks, demonstrating its superiority for virus evolutionary drivers prediction.

### 2.3 E2VD captures fundamental patterns of virus evolution

First, we conduct module ablation studies to explore the contributions of the **L**ocal-**G**lobal dependence coupling (LG) module and **M**ulti-**T**ask focal learning (MT) module to prediction performance. As shown in Fig.4a, four sets of experiments are designed for comparison, including E2VD (referred to as Ours), E2VD without the LG module (referred to as w/o LG), E2VD without the MT module (referred to as w/o MT), and E2VD without both modules (referred to as w/o MT&LG). Compared with the E2VD without both modules, the introduction of the LG module is beneficial to both classification and regression tasks, with the performance of the regression task being significantly improved (PCC: 0.385 to 0.871). In contrast, the MT module significantly improves the performance of classification task (Accuracy: 50% to 68.52%). It is worth noting that the Recall is sharply improved (Recall: 0 to 69.63%), which means that MT is effective in mining rare beneficial mutations for virus fitness. The combination of the above two modules further improves the performance of both classification and regression tasks, reaching the Accuracy of 91.11%, the Recall of 96.3%, and the PCC of 0.87. We also conduct the same module ablation experiments with ESM-2 (650M) and ProtT5 (3B) as the extractor for sequence embeddings (Supplementary information, Table.S11, Table.S12), and the results are consistent with the above conclusions.

**Fig. 4.**
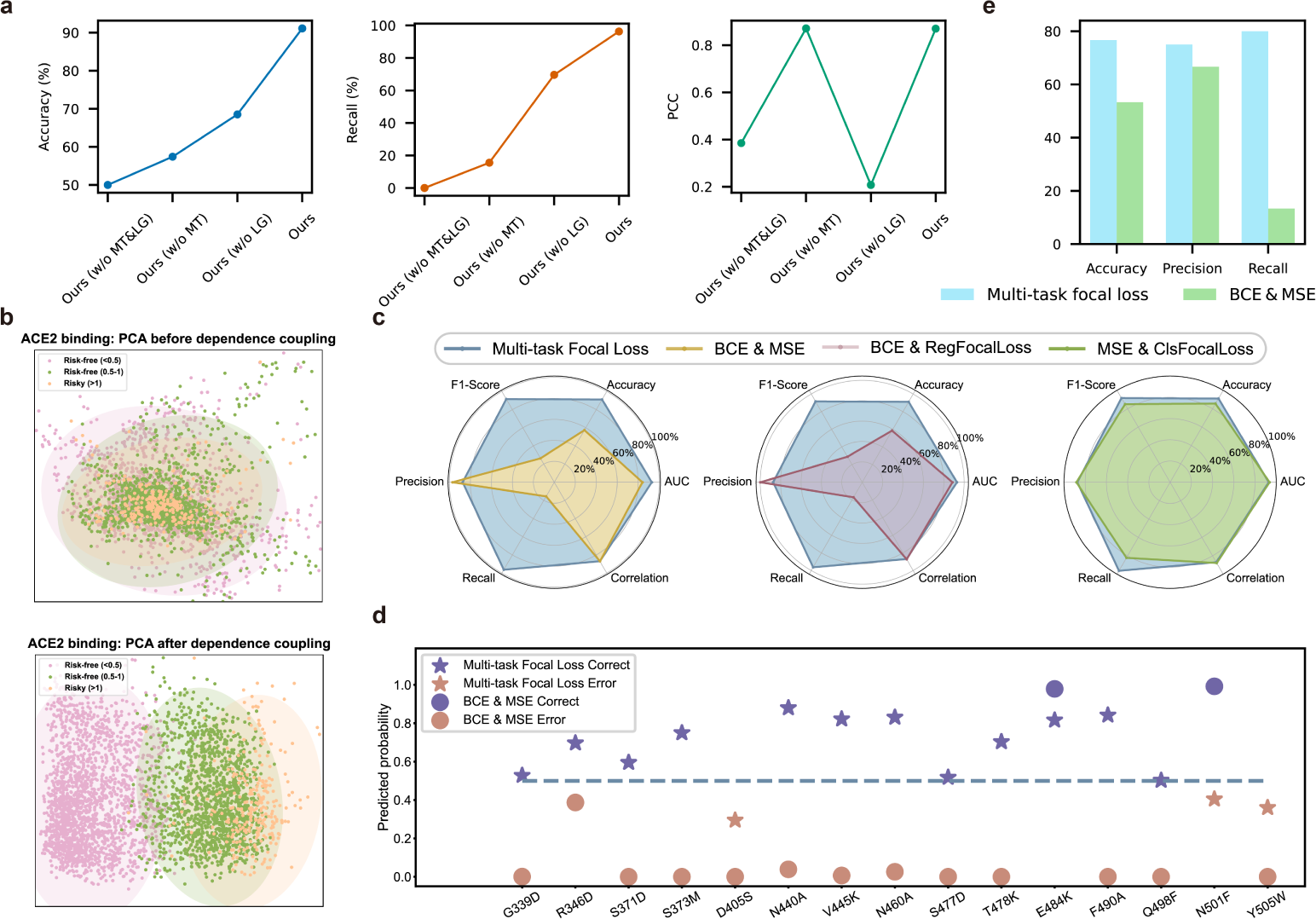
Ablation studies. **a**, Module ablation studies. LG refers to local-global dependence coupling module, and MT refers to multi-task focal learning module. **b**, Dimensionality reduction visualization of different types of mutations of the binding affinity prediction task. Three types of mutations are presented, including Risk-free (*Ka ratio<*0.5), Risk-free (0.5*<Ka ratio<*1), and Risky (*Ka ratio>*1). **c**, The ablation studies on multi-task focal loss to analyze the impact of single focal loss on prediction performance. **d**, Prediction performance of our proposed multi-task focal loss and BCE&MSE on beneficial mutations. **e**, The overall prediction performance comparison of multi-task focal loss and BCE&MSE on the comprehensive dataset consisting of both beneficial and harmful mutations.

Second, we perform dimensionality reduction visualization of different types of mutations using Principal Component Analysis (PCA), as shown in Fig.4b. The three types of mutations described by risk level (Methods 4.9) of binding affinity prediction task cannot be distinguished in the feature space before being processed by the LG module, showing a very high degree of overlap. Upon processing by the LG module, features of different mutations are distinctly separated with clear boundaries. It can be inferred that LG enhances E2VD’s sensitivity to various mutation types by restoring the intramolecular interaction system, enabling a better understanding of the evolutionary fitness of the virus. We also perform dimensionality reduction visualization on the datasets of the expression and antibody escape prediction task, and the results further support the above conclusion (Supplementary information, Fig.S1).

Third, we conduct ablation experiments on multi-task focal loss to analyze the impact of single focal loss on prediction performance. As shown in Fig.4c, we compare four different loss functions for multi-task learning, including the combination of Mean Squared Error (MSE) loss and Binary Cross Entropy (BCE) loss, BCE with focal loss for regression (referred to as RegFocalLoss), MSE with focal loss for classification (referred to as ClsFocalLoss), and the proposed multi-task focal loss. We find that the introduction of RegFocalLoss or ClsFocalLoss alone improves the overall prediction performance, and the full multi-task focal loss further improves the overall performance, with the Accuracy increasing from 57.41% to 91.11% and the Recall increasing from 15.56% to 96.30%.

Fourth, in order to assess the ability of multi-task focal loss to capture rare beneficial mutations, we randomly select 15 beneficial (associated with increased binding affinity) and 15 harmful (associated with reduced binding affinity) mutations to form the test set (Methods 4.10). As shown in Fig.4d, E2VD with multi-task focal loss achieves excellent prediction performance on beneficial mutations, correctly predicting 12 out of 15 beneficial (risky for humans) mutations, i.e. 80% of the total. In contrast, the prediction performance of E2VD with BCE&MSE lags far behind, only correctly predicting 2 mutations, i.e. about 13% of the total. For harmful mutations, multi-task focal loss and BCE&MSE achieve similar performance, correctly predicting 11 and 14 respectively (Supplementary information, Fig.S2). As we expected, since BCE&MSE cannot help the model learn rare beneficial mutations, the model tends to predict all mutations as harmful (14 out of 15). Considering both beneficial and harmful mutations in Fig.4e, three indicators, including Accuracy, Precision, and Recall, are adopted to evaluate overall prediction performance. Multi-task focal loss outperforms the BCE&MSE comprehensively and significantly, with a Recall improvement of more than 6 times.

In summary, E2VD sensitively distinguishes different types of mutations by amplifying the subtle effects caused by few mutations through local-global dependence coupling module, and accurately identifies rare risky (i.e. beneficial) mutations through the hard sample mining strategy of multi-task focal learning module, thereby capturing the fundamental patterns of virus evolution.

### 2.4 Generalization performance of E2VD across different lineages

Viruses continue to evolve under selective pressure, resulting in the development of various lineages of SARS-CoV-2 [52]. Therefore, the generalizability of virus variation drivers predictor is crucial to cope with intricate evolutionary trends. Considering that the wet-lab experiment data used for training is severely affected by batch effects [53], it becomes challenging to predict precise values of variation drivers, thereby hindering the efficiency of identifying high-risk variants. On the contrary, predicting the relative order can help sort variants according to evolutionary fitness, providing more valuable guidance. Inspired by the fact that inverse pairs can reflect the degree of disorder of the sequence [54, 55], we propose **O**rdinal **P**air **P**roportion (OPP) to evaluate the generalization performance of different models. As shown in Fig.5a, a correctly predicted pair is defined as two mutations whose order in the predicted fitness landscape matches that of the ground truth fitness landscape. Otherwise, it is considered as a mis-predicted pair, i.e. an inverse pair. OPP represents the proportion of correctly predicted mutation pairs in all mutation pairs, reflecting the degree of disorder of the predicted fitness landscape. The larger the value of OPP, the less chaotic the predicted fitness landscape, showing that the model is suitable for predicting the relative order of virus variation drivers.

**Fig. 5.**
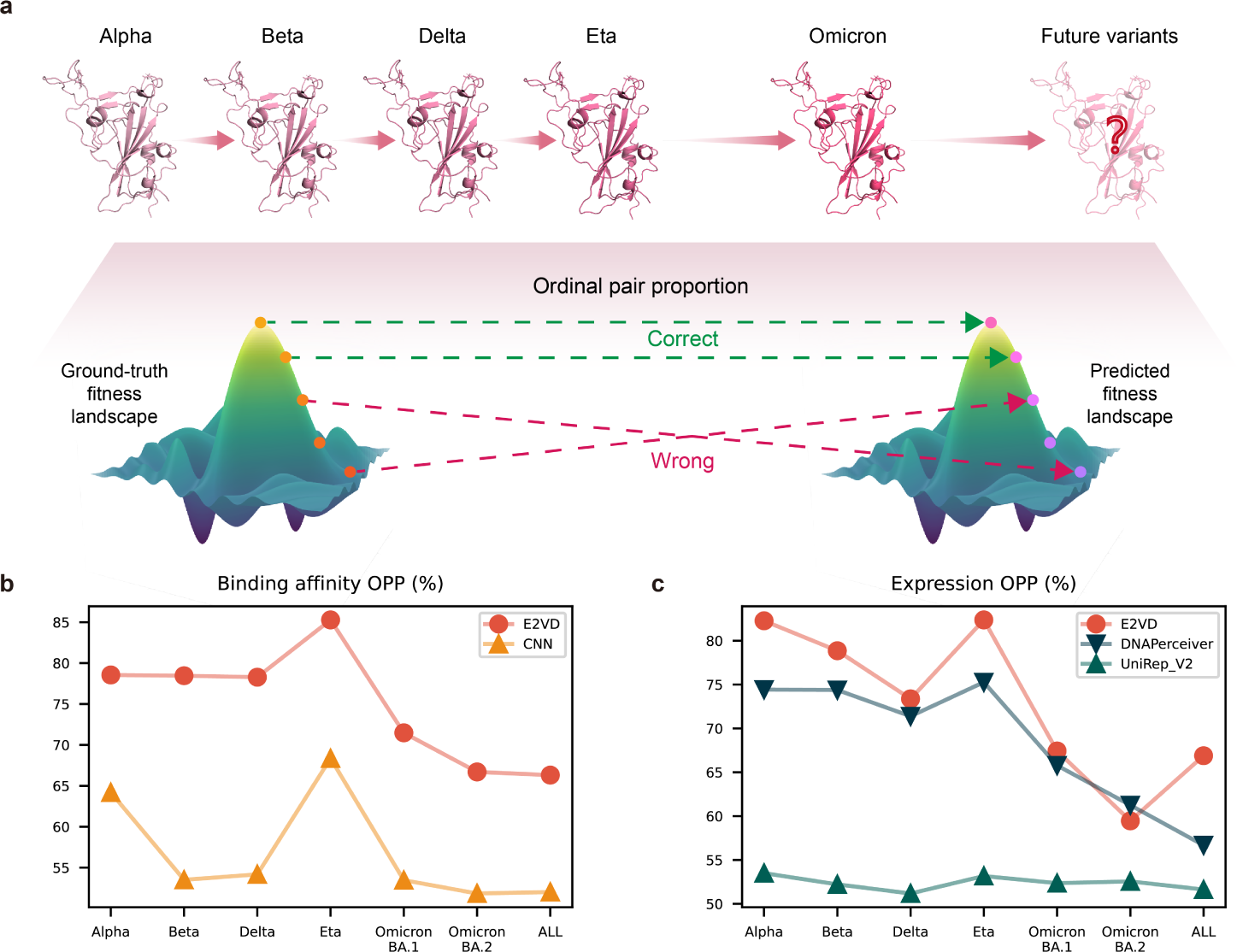
Generalization performance evaluation. **a**, The proposed **O**rdinal **P**air **P**roportion (OPP) indicator for the generalization performance evaluation of different models. It represents the proportion of correctly predicted pairwise mutations in all mutation combinations, reflecting the degree of chaos of the predicted fitness landscape. **bc**, OPPs of different models on the binding affinity (b) and expression prediction tasks (c). Six different lineages as well as a mixture of all lineages (referred to as All) are adopted to evaluate the generalization performance.

With the proposed OPP metric, we assess the generalization capabilities of different models on virus variation drivers prediction tasks. We transfer the model trained with deep mutational scanning (DMS) dataset of wild-type RBD [12] to perform zero-shot inference on different lineages [13]. As different lineages contain multiple mutations relative to the wild-type, this can be considered a task of out-of-distribution generalization [56]. It is worth noting that we conduct generalization assessment on virus variation drivers prediction tasks except for antibody escape task. This is because the DMS dataset of antibody escape [14] only involves the wild-type RBD, resulting in a lack of data of other lineages.

As shown in Fig.5b, in the binding affinity prediction task, OPPs of CNN and E2VD are calculated for six different lineages as well as a mixture of all lineages (referred to as All) to evaluate the generalization performance. We can see that E2VD comprehensively and significantly surpasses CNN, whether on a single lineage or the mixture of all lineages. Specifically, E2VD’s prediction performance remains stable above 78% on the four lineages (Alpha, Beta, Delta, Eta) before Omicron (B.1.1.529), but there is a clear performance decay on the two sub-lineages (BA.1, BA.2) of Omicron. Lineages before Omicron contain only few mutations in RBD, while Omicron contains 15 mutations in RBD [49]. This makes it more difficult to generalize to the Omicron sub-lineages, leading to significant decrease in prediction accuracy. Therefore, Omicron is a reasonable dividing line for model updating, that is, if we predict the virus variation drivers of the lineages after Omicron, retraining the model with the DMS datasets of the lineages that appears after Omicron is necessary. Taking all lineages together, CNN’s OPP is improved by 14.3% by E2VD. The situation is similar in expression prediction task (Fig.5c), where E2VD outperforms the other two competitors on most lineages, with the OPP achieving an 10.2% improvement over DNAPerceiver [24] for the mixture of all lineages. Overall, E2VD surpasses state-of-the-art methods across the board on out-of-distribution lineages, demonstrating enhanced generalization performance.

### 2.5 E2VD perceives evolutionary trends

E2VD has shown superior prediction performance and generalization in multiple virus variation drivers prediction tasks, making it possible to predict viral evolutionary trends. In this work, we demonstrate a flexible combination pipeline that assembles *in silico* deep mutational scanning with three different variation drivers predictors developed by E2VD for viral evolutionary trends prediction, as shown in Fig.6a. Specifically, referring to the practice of wet-lab deep mutational scanning [12, 57], we first sequentially mutate a single-site amino acid in the RBD to other 19 types, and complete the *in silico* deep mutational scanning in the entire region site by site. Then, the binding affinity, expression, and antibody escape induced by each single-site mutation are predicted by the corresponding predictors for subsequent analysis.

**Fig. 6.**
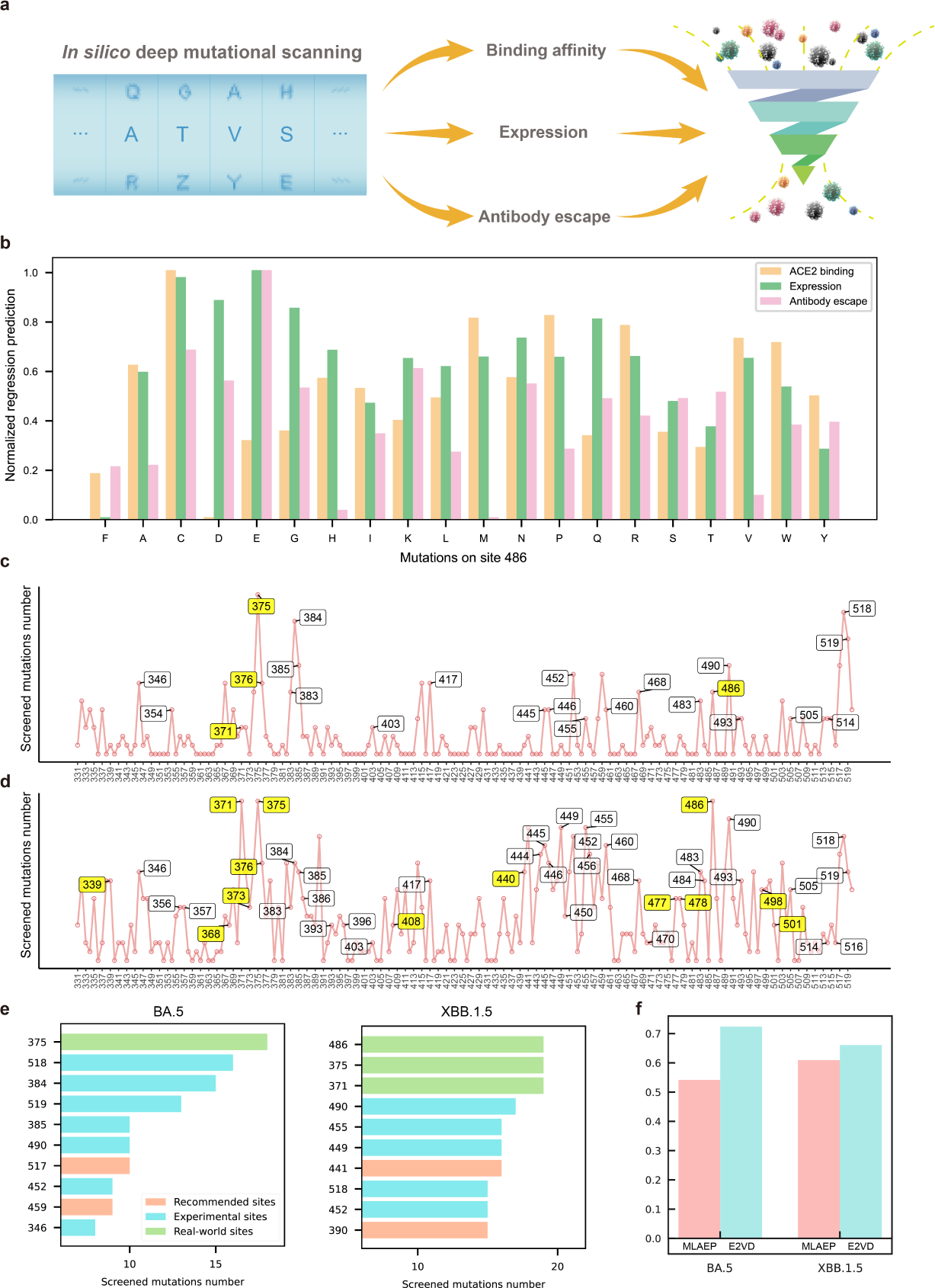
The pipeline and results of evolutionary trends prediction. **a**, A flexible combination pipeline that assembles *in silico* deep mutational scanning with three different variation drivers predictors for virus evolutionary trends prediction. **b**, The normalized regression predictions of the three key variation drivers (binding affinity, expression, and antibody escape) after *in silico* deep mutational scanning on 486 site of XBB.1. **cd**, The distribution of the mutational risk of each site of BA.5 (c) and XBB.1.5 (d), where the “screened mutations number” represents the remaining mutations at a site after screening. The sites marked in yellow boxes are the risky ones that have been mutated after BA.5 or XBB.1.5 lineage in the real world, and the sites marked in white boxes are the risky ones recommended by wet-lab experiments. **e**, The top 10 recommended mutation sites in BA.5 (left) or XBB.1.5 (right), in which the hit rate (real-world mutation sites and wet-lab experimental recommended sites) both reach 80%. **f**, Area Under Curve (AUC) of E2VD and the competitor on BA.5 and XBB.1.5 lineages in the high-risk mutation site prediction task.

First, we demonstrate the ability of our E2VD-core pipeline to reveal the intrapandemic evolution caused by viral fitness changes. XBB.1.5 showed a significant growth advantage over XBB.1 during its rapid rise to become the dominant lineage [58, 59], even though they only differed by one S486P mutation. In order to explore the essential connection between this mutation and transmissibility, we perform *in silico* deep mutational scanning on 486 site of XBB.1 and subsequently predict the three key variation drivers (binding affinity, expression, and antibody escape) for all mutations (Fig.5b). We find that when the Ser on 486 site in XBB.1 is mutated to Pro (i.e. XBB.1.5), its binding affinity with human ACE2 surges and its expression level is also improved to a certain extent. Besides, antibody escape exhibits a slight weakening. The prediction results above are basically consistent with previous wet-lab measurements [12, 13, 58], allowing us to draw the following conclusion. XBB.1.5 achieves a surge in binding affinity while maintaining qualified antibody escape, which likely contributes to its significant growth advantage.

Second, we demonstrate how our E2VD-core pipeline recommends potential high-risk mutation sites of a lineage. After performing the *in silico* deep mutational scanning for each site in a lineage, we screen the created single-mutation variants by setting different thresholds of predicted biological drivers according to their importance (Methods 4.12.3). The distributions of the mutational risk of each site of BA.5 and XBB.1.5 are shown in Fig.6cd, where the “screened mutations number” represents the remaining mutations (i.e., single-mutation variants) at a site after screening. The higher the number of remaining mutations at a site, the higher the risk of this site. Thus, our pipeline recommends the most risky mutation sites based on the number of screened mutations of each site. As shown in Fig.6e, for the top 10 recommended mutation sites in BA.5 or XBB.1.5 lineage, the hit rate (real-world mutation sites and wet-lab experiment recommended sites) both reach 80%, proving the ability of our pipeline to accurately recommend high-risk mutation sites. Furthermore, taking the wet-lab deep mutational scanning statistic results [14, 15] as ground-truth, we plot the Receiver Operating Characteristic (ROC) curves for the two lineages and calculate the corresponding Area Under Curve (AUC) (Fig.6f). We found that E2VD-core pipeline comprehensively surpasses the competitor MLAEP-core pipeline [17] on two lineages, where improves the AUC on BA.5 by 18.2% and that on XBB.1.5 by 5.2%.

In summary, the flexible combination pipeline not only reveals intrapandemic evolution caused by viral fitness changes, but also accurately recommends high-risk mutation sites in different lineages, demonstrating the perception of virus evolutionary trends.

## 3 Discussion

E2VD provides an alternative to costly wet-lab experimental approaches for the analysis and prediction of virus evolutionary fitness in a unified and interpretable way, holding great potential for enhancing the rapid response to emerging viral infections. It should be emphasized that SARS-CoV-2 is studied as a typical case in this work due to the extensive availability of deep mutational scanning data for it. However, E2VD is not limited to specific viruses. The “few-site mutations” and “rare beneficial mutations” are general characteristics faced by all viruses in the process of evolution. Therefore, our evolution-driven framework designed on this basis can potentially be extended to a variety of viruses such as HIV, MERS-CoV when there are enough deep mutational scanning data for training.

E2VD currently faces certain limitations, such as the lack of support for predicting mutations of non-equal lengths (amino acid insertions or deletions) and a significant decrease in predictive accuracy when applied to Omicron variants. Existing deep mutational scanning data of viral proteins typically only include amino acid substitutions, that is, data on amino acid insertions or deletions are very scarce, which makes it difficult to train deep learning models that support non-equal length mutation prediction. with the increasing amount of deep mutational scanning data of non-equal length mutations of viral proteins, it is promising to train qualified models for the prediction of few-site mutation impacts caused by amino acid insertions or deletions. Facing the future, we can further improve model generalization performance to promote the expansion of the effectiveness boundary from both data and model architecture. With the growing availability of deep mutational scanning data from specific lineages, incorporating them into the training dataset can effectively improve the generalization of the trained model. Besides, the reasonable introduction of modules such as domain adaptation [60] is also expected to help the model generalize better. Last but not least, in this work, we predict the mutational trends of viruses based on the evolutionary fitness of viral proteins, without integrating high-level viral characteristics directly, including pathogenicity and transmissibility, etc. Incorporating the above-mentioned high-level viral characteristics into our framework is a potential next step to connect selective pressures within and outside the host.

## 4 Methods

### 4.1 Data

#### 4.1.1 Benchmark dataset of binding affinity prediction task

For binding affinity prediction task, we adopt the deep mutational scanning (DMS) dataset of binding affinity from previous study [12]. The above work reported the difference in log10*K_d_*, i.e. apparent dissociation constant, relative to wild-type SARS-CoV-2 RBD. The raw data form is that one mutation corresponds to one Δ log_10_ *K_d_*, and we preprocess the raw dataset to facilitate model training. First, based on the mutations, we substitute residues at corresponding positions on wild-type RBD sequence to obtain the mutated RBD sequences. Second, we convert Δ log_10_ *K_d_* to *K_a_ ratio* to get the training labels representing the binding strength as follows:

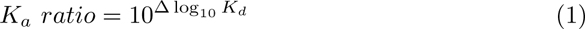

where *K_a_ ratio* represents the ratio of the binding strength with human ACE2 between the mutated RBD and the wild-type RBD. A value greater than 1 represents an increase in binding affinity, and a value less than 1 represents a decrease in binding affinity.

Finally, there are 3,702 samples, including 3,388 negative samples (harmful for fitness) with label less than 1, and 314 positive samples (beneficial for fitness) with label greater than 1. To evaluate the performance of E2VD on binding affinity task, 5-fold cross-validation training is adopted. The blind test set (n=54) keeps the same as previous publication [19], which includes the same number of positive and negative samples. The prediction performance on the blind test set is calculated as the mean of the metrics of five trained models. The validation set (n=110) of each fold randomly samples the same number of positive and negative samples to avoid the imbalance of sample categories affecting the verification results, and the remaining samples form the training set (n=3,538) of each fold.

#### 4.1.2 Benchmark dataset of expression prediction task

For expression prediction task, we adopt the DMS dataset of expression from previous study [12]. This work represented expression as Δ log(*MFI*) relative to wild-type SARS-CoV-2 RBD. Similarly, we substitute residues at corresponding positions on wild-type RBD sequence to obtain the mutated RBD sequences, and calculate the training label as follows:

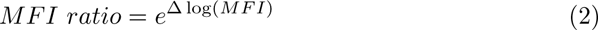

where *MFI ratio* represents the ratio of the expression level of the mutated RBDs relative to the wild-type RBD. A value greater than 1 represents an increase in expression, and a value less than 1 represents a decrease in expression.

Finally, there are 3,781 samples, including 436 positive samples (beneficial for fitness) with labels greater than 1 and 3,345 negative samples (harmful for fitness) with labels less than 1. To evaluate the performance of E2VD on expression prediction task, 5-fold cross-validation training is adopted. The blind test set (n=100) randomly sample the same number of positive and negative samples, and the average of metrics of five trained models is considered as the performance result. The validation set (n=120) of each fold randomly samples the same number of positive and negative samples, and the remaining samples form the training set (n=3,561) of each fold.

#### 4.1.3 Benchmark dataset of antibody escape prediction task

For antibody escape prediction task, we adopt the DMS dataset of wild-type RBD with different antibodies from previous study [14]. This work conducted deep mutational scanning between 3,051 antibodies and the wild-type RBD, reporting antibody names, mutations on wild-type RBD, and the corresponding escape scores. In this work, we use the escape scores as training labels directly. According to the previous publication [61], 0.4 is a relatively high escape score, so we assume that it is an appropriate threshold to differentiate between high and low antibody escape risk. Taking 0.4 as the threshold of escape risk, samples with scores greater than 0.4 are considered to have a high escape risk and the rest are considered to have low escape risk.

Finally, there are 57,740 positive samples (high risk) and 284,032 negative samples (low risk) in the dataset. To evaluate the performance of E2VD on antibody escape task, we randomly sample the same number of positive and negative samples as test set (n=9,600) and the remaining samples form the training set (n=332,172).

### 4.2 Protein language model customization

We adpot UniRef90 dataset consisting of 144 million raw protein sequences as pre-training dataset. Unlike the Uniref100 dataset, the Uniref90 dataset offers rich evolutionary information without compromising performance in the early training stages [41]. This richness aids in enhancing zero-shot generalization by exposing the protein language model to a diverse range of sequence examples from various protein families. The architecture of customized protein language model is similar to *BERT_LARGE_* [62], and it is pre-trained with only the objective of masked language modeling with the original masking probability of 15%. We follow the widely recognized configurations and two-stage pre-training strategy of mainstream protein language models ProtTrans [40] and natural language models *BERT* [63]. The efficiency gain of two-stage pretraining strategy comes from optimizing resource utilization using shorter sequences, thereby speeding up the convergence of longer sequences [40, 63]. During preprocessing, non-generic or unresolved amino acids are mapped to unknown (represented by X). In the first pre-training stage, the raw protein sequences are truncated with a length of 128, resulting in a training dataset of 459.3 million fragments. The protein language model is pre-trained with a learning rate of 0.0004, warm-up steps of 40,000, and epochs of 65. In the second pre-training stage, the raw protein sequences are truncated with a length of 256, resulting in a training dataset of 265.5 million fragments. The protein language model undergoes continuous pre-training with a learning rate of 0.0001 and epochs of 35. It is worth noting that we retain all truncated protein fragments for pre-training during the sequence truncation process, that is, no fragments are discarded. During pre-training, the Lamb optimizer and weight decay of 0.01 are adopted as ProtBert [40]. The detailed configuration is as follows: 24 layers, 1,024 hidden dimension, 16 attention heads, 64 local batch size, and 16,384 global batch size.

### 4.3 Architecture overview

The architecture of E2VD includes three parts: protein language model, local-global dependence coupling and multi-task focal learning. The local-global dependence coupling module consists of CNN for establishing local dependencies, and ContextPool-attention for establishing global dependencies. In the multi-task focal learning module, our model performs multi-task learning including classification and regression tasks under the supervision of the proposed multi-task focal loss.

Specifically, we denote as protein language model, as local dependence establishing, as global dependence establishing, as focal learning under classification task, and as focal learning under regression task. Given an input protein sequence *S*, the protein language model extracts the residue-level features *E* of *S* as *E* = (*S*). The extracted features are further refined by local-global dependence coupling module, concatenated and pooled into sequence-level feature *F* as follows:

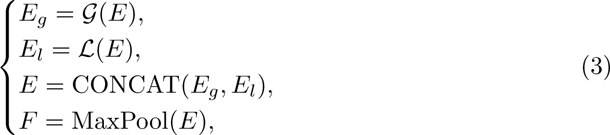

where *E* denotes residue-level features, *E_g_* and *E_l_* denote global and local residue-level features, *F* denotes sequence-level features, CONCAT represents concatenate, and MaxPool represents max pooling.

Subsequently, the focal learning under classification task predicts whether the variation drivers increase or not, and the focal learning under regression task predicts the specific values of variation drivers.

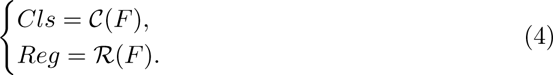

where *Cls* denotes the output of classification task, *Reg* denotes the output of regression task.

### 4.4 Sequence tokenizing and embedding

An input protein sequence is treated as a sequence of tokens (residues), and the tokenizer maps every token to its corresponding index. For protein sequences with different lengths, the tokenizer uses the reserved token ‘[PAD]’ to fill the sequence to the longest, and set the value of the corresponding attention mask to 0. The features of amino acids are extracted in the form of embeddings, i.e., vector representations. Specifically, let *l* be the longest length of sequences and *d* be the dimension of the embedding. Embeddings extracted by different protein language models have different dimension *d*. For our customized protein language model and ProtT5 (3B), *d* are 1024. For ESM2-650M, ESM2-3B and ESM2-15B, *d* are 1280, 2560 and 5120, respectively. The length of the tokenized sequence is *l*, and the shape of the embedding for each sequence is *d × l*. After extracting embeddings, we conduct standardization to enlarge the difference between different sequences. In order to distinguish the impact of standardization and our proposed local-global dependence coupling module on prediction performance, we perform ablation experiments (Supplementary information, Table.S8).

Specifically, let *X_d×l_* be the extracted embeddings, and *N* be the number of sequences. First, the embeddings are flattened into one dimension 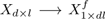. For feature dimension *i ∈* [1, *dl*], the standardization is as follows:

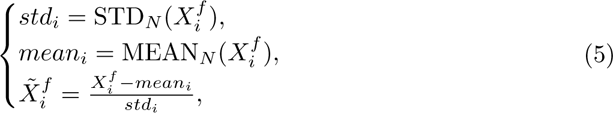

where STD*_N_* (·) and MEAN*_N_* (·) denote the standard deviation calculation and the mean calculation among all the *N* embeddings respectively. After standardization, the flattened embedding 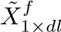 is reshaped into the original dimension 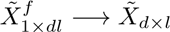.

For binding affinity and expression prediction, the input is only the mutated RBD. For antibody escape prediction, the input involves RBD and antibody sequences. Considering that the antibody variable region includes heavy and light chains, we conduct standardization separately to the extracted embeddings of RBD, antibody heavy chain and antibody light chain as described above.

### 4.5 Local-global dependence coupling

We use ContextPool-attention and CNN to capture the global dependencies and the local dependencies of extracted embeddings, respectively.

#### 4.5.1 ContextPool-attention

Context pooling [45] adopts an explicit way to learn context-aware token features by learning to pool neighboring features for each token. After that, self-attention between such pooled features can thus be context-aware and model high-level dependencies. Inspired by this, we set the learnable granularity to establish global dependencies of mutations based on seeking for motifs in protein sequences.

Specifically, a mapping function *m*(·) is learned to predict both the pooling weights *w* ∈ R*^n×1^* and sizes *s* ∈ R*^n×1^* for *l* input token features 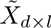, i.e. 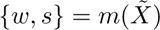. Here, we implement *m*(·) using three convolutional layers with softmax function behind, and the output channel size of the last layer is set to 2 to generate the vectors *w* and *s* simultaneously. The normalized pooling size *s_i_* is transformed into standard deviation of Gaussian mask 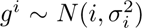, where *σ_i_* = *rn ∗ s_i_*, and *r* is a scalar empirically set as 0.1.

The final context pooling function multiplies the learned pooling weights *w* with Gaussian mask *g^i^* for token 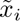. Specifically, the context pooling function is calculated as follows:

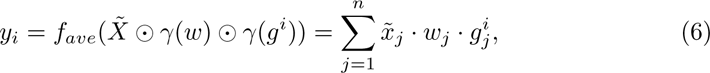

where *y_i_∈ Y* represents the context pooling features, *f_ave_* denotes average pooling function, and *γ*(*·*) is broadcasting function for element-wise multiplication *⊙*.

#### 4.5.2 CNN

We adopt a one-dimension convolution neural network to establish the local dependencies between the mutation and its neighboring residues. The CNN here consists of 3 convolution layers with Layer Normalization and uses leaky rectified linear unit (Leaky ReLU) as activation function. We set the kernel size to 3 and the stride to 1 to ensure the shape of the output is the same as the input.

#### 4.5.3 Feature coupling

The outputs of ontextPool-attention and CNN are concatenated together to get the final extracted features of residues, and the feature of the whole sequence is calculated through max pooling on the residue dimension. It is worth noting that, for antibody escape prediction, the embeddings of RBD, antibody heavy chain, and antibody light chain are input into independent local-global dependence coupling channel respectively to capture corresponding global dependencies and local dependencies. The features of each kinds of sequence are calculated through max pooling on the residue dimension, and the results are concatenated as the final feature to predict antibody escape.

### 4.6 Multi-task focal learning

We propose multi-task focal learning module to alleviate the imbalance between positive and negative samples through multi-task learning manner, in which the fitting strength to hard-to-learn samples is increased. We adopt two focal losses for different training tasks, i.e. the focal loss for classification and the focal loss for regression.

For classification task, the training intensity of a certain category can be adjusted by assigning different weights to samples with different labels in focal loss for classification. Specifically, the calculation of the focal loss for classification is as follows:

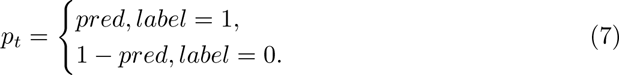

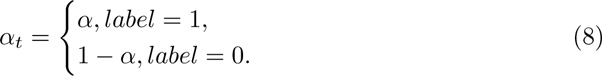

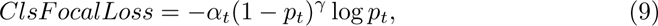

where *pred* stands for the prediction of the classification model, *label* stands for the classification label. The penalty index *γ* indicates the punishment intensity for large error samples. The larger *γ* is, the lower the tolerance to wrong samples will be. The weight coefficient *α* can adjust the attention of loss function to positive and negative samples.

For regression task, the calculation of the focal loss is as follows.

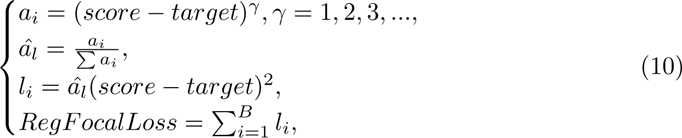

where *score* stands for the prediction of the model, *target* stands for the regression label, and 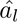 is the scaled error in case the gradient value is too small. The penalty index *γ* is the same as that in the focal loss for classification.

The final multi-task focal loss function is as follows:

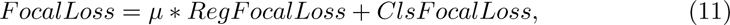

where *µ* denotes the weight of the regression loss. We set *µ* = 1 in this work.

### 4.7 Evaluation metrics

#### 4.7.1 Metrics for regression tasks

For regression tasks, we use Mean Squared Error (MSE) and Pearson’s Correlation Coefficient (PCC). Specifically, let *y_pred_* be the predicted *K_a_ ratio* and *y_label_*be the experimental *K_a_ ratio*, the MSE and PCC are calculated as follows

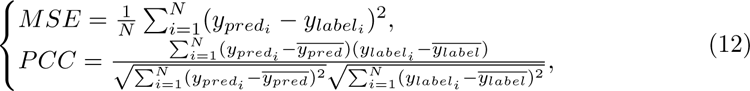

#### 4.7.2 Metrics for classification tasks

The metrics for classification tasks includes *AUC*, *Accuracy*, *F* – 1 *Score*, *Precision* and *Recall* between label and prediction.

*AUC*, i.e. *AU – ROC*, indicates the probability that the predicted score of positive samples is higher than the one of negative samples. Specifically, let *m*^+^ and *m^−^* be the number of positive and negative samples respectively. *D*^+^ and *D^−^* denote the set of positive and negative samples respectively. *AUC* is calculated as follows:

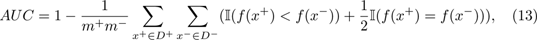

where the *f* (·) denotes the prediction function, and I(·) values 1 if the input is *True*, otherwise it values 0.

For binary classification tasks, the predictions can be divided into four cases: *TP* if the prediction and label are both positive, *TN* if the prediction and label are both negative, *FP* if the prediction is positive and the label is negative, *FN* if the prediction is negative and the label is positive. Therefore, the other four metrics are calculated as follows:

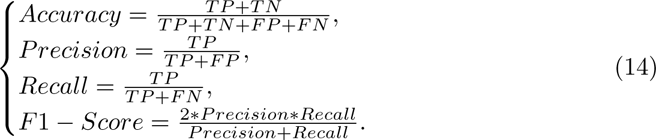

#### 4.7.3 Ordinal Pair Proportion

The inverse pairs can reflect the degree of disorder of the sequence, in which the greater the number of inverse pairs in a sequence, the higher the degree of disorder of the sequence. If the number of inverse pairs is small, the sequence is relatively ordered. Inspired by this, we propose a new metric, *Ordinal Pair Proportion*(*OPP*), to evaluate generalization performance of different models. Specifically, we first combine all the prediction results into binary pairs. For each pair, if the label order of the two predictions is consistent with the order of prediction, we call the pair *ordinal*, and the *OPP* is the proportion of the *ordinal* pairs in all pairs. Then, for the prediction set *N, n* = *|N |*, *OPP* is calculated as follows:

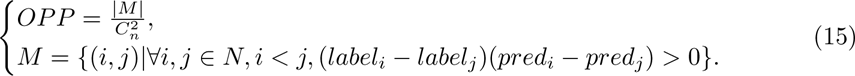

### 4.8 Ablation studies

We conduct ablation studies to explore the individual contributions of different components of the model to prediction performance, including protein language model ablation study, module ablation study, and loss function ablation study.

#### 4.8.1 Protein language model ablation experiments

To evaluate the effectiveness of customized pre-training configuration on the prediction performance, we compare it with well-known pre-trained PLMs. Specifically, ProtT5 (3B) in the ProtTrans series [40], which achieved excellent performance at both the residue-level and protein-level, is selected. Among the ESM series, ESM-1v (650M) [41], which has excellent zero-shot prediction ability, and ESM-2 (15B) [29] with the largest parameter size, are selected. In particular, due to terrible memory usage and training time, we adopt ESM-2 (650M) rather than ESM-2 (15B) in antibody escape prediction task.

#### 4.8.2 Module ablation experiments

We conduct ablation experiments on local-global dependence coupling module and multi-task focal learning module with three different protein language models to demonstrate the generalizability of the results, including ESM-2(650M), ProtT5(3B) and our customized protein language model. The benchmark dataset of binding affinity prediction task is adopted and the experiment results can be found in Supplementary information (Table.S10,Table.S11, and Table.S12).

#### 4.8.3 Loss ablation experiments

We conduct ablation experiments on multi-task focal loss to analyze the impact of single focal loss on prediction performance. Specifically, we adopt the benchmark dataset of binding affinity prediction task and set four different combinations of loss functions, including binary cross entropy loss paired with mean squared error loss, binary cross entropy loss paired with focal loss for regression, mean squared error loss paired with focal loss for classification, and the multi-task focal loss. The experiment results can be found in Supplementary information (Table.S13).

### 4.9 Dimensionality reduction visualization of different types of mutations

Mutations that are beneficial to the evolutionary fitness of the virus are risky to humans, which we call “risky mutations”. Similarly, mutations that are harmful to the evolutionary fitness of the virus are risk-free to humans, which we call “risk-free mutations”. In three different downstream tasks, the above “risky mutations” and “risk-free mutations” are defined by different indicators. First, for binding affinity prediction task, the fitness of the viral protein caused by the mutations is measured by *K_a_ ratio*, which represents the ratio of the binding strength with human ACE2 between the mutated RBD and the wild-type RBD. Considering that risk-free mutations (i.e. the harmful mutations for virus) account for the majority in the deep mutational scanning (DMS) dataset, we can further divide this type of mutations into two subtypes with 0.5 as the threshold. Therefore, three types of mutations are defined in binding affinity prediction task, including Risk-free (*K_a_ ratio<*0.5), Risk-free (0.5<*K_a_ ratio<*1), and Risky (*K_a_ ratio>*1). Second, for the expression prediction task, the fitness of the viral protein caused by the mutations is measured by *MFI ratio*, which represents the ratio of the expression level of the mutated RBDs relative to that of the wild-type. Therefore, the definition of mutation types is similar to that of binding affinity prediction task, in which three types of mutations are defined, including Risk-free (*MFI ratio <*0.5), Risk-free (0.5*<MFI ratio<*1), and Risky (*MFI ratio>*1). Third, for the antibody escape prediction task, the fitness of the viral protein caused by the mutations is measured by escape score of DMS data. Taking 0.4 as the threshold of escape risk, samples with scores greater than 0.4 are considered to have a high escape risk and the rest are considered to have low escape risk. Therefore, different from binding affinity and expression prediction task, two types of mutations are defined here, including Low-risk (*Escape score<*0.4) and High-risk (*Escape score>*0.4).

We adopt Principal Component Analysis (PCA) for feature dimensionality reduction, in which the features before and after local-global dependence coupling module are visualized. For binding affinity and expression prediction tasks, we choose the RBD embedding from protein language model as the feature before dependence coupling. For antibody escape prediction task, we concatenate the RBD embedding, antibody heavy chain embedding and antibody light chain embedding from protein language model as the feature before dependence coupling. The feature that passes through local-global dependence coupling module are used as the features after dependence coupling. Finally, we color the samples according to the their risk level.

### 4.10 Experiments to mine rare beneficial mutations

We select XBB.1.5 as the reference lineage, and randomly select 15 mutations with increased binding affinity on real-world mutation sites within its RBD region ^1^ from DMS dataset. Next, we randomly select 15 mutations with reduced binding affinity at each beneficial mutation site to form a test set with balanced positive and negative samples to ensure comprehensiveness of the evaluation. Taking remaining samples as training data, we train two models with multi-task focal loss and BCE&MSE loss respectively and compare their prediction performance.

### 4.11 Generalization performance experiments

We calculate *Ordinal Pair Proportion*(*OPP*) metrics on binding affinity and expression prediction tasks to evaluate the generalization performance of different models. All models are trained using DMS data of wild-type RBD [12] and subsequently perform zero-shot inference on DMS data of multiple lineages [13].

We calculate OPP metrics on all samples from different lineages, as well as all samples within each lineage, including Alpha, Beta, Delta, Eta, Omicron BA.1 and Omicron BA.2. Specifically, samples within each lineage *V* is denoted as *Seq_V_*. The samples of multiple lineages datasets *Seq* includes *Seq* = *Seq_Alpha_, Seq_Beta_, Seq_Delta_, Seq_Eta_, Seq_Omicron BA_*_1_, *Seq_Omicron BA_*_2_.

The *OPP* of all samples from different lineages can be calculated as follows:

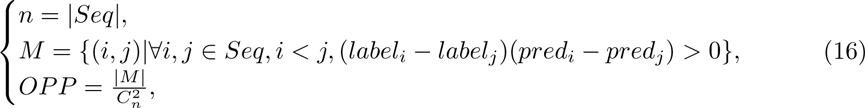

where *pred_i_* and *label_i_* are the predictions and labels of binding affinity and protein expression respectively.

The *OPP* of all samples within each lineage *V* can be calculated as follows:

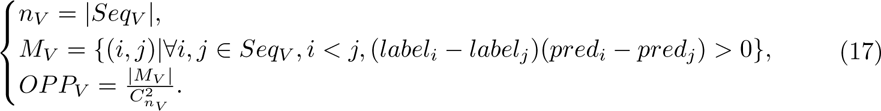

### 4.12 Evolutionary trends prediction experiments

#### 4.12.1 Model updating

Since the selected lineages to be studied here are all branches of Omicron, the DMS datasets of the lineages emerging after Omicron are selected for model retraining to avoid a significant decrease in model generalization caused by the notable difference in lineages before and after Omicron.

The binding affinity predictor is retrained with the DMS dataset of Omicron BA.1 and Omicron BA.2 [13]. We normalize the provided log_10_ *K_d_* values with that of Omicron BA.2 to get the training labels. Samples with labels that are greater than 1 are considered as positive samples, while those with labels that are less than 1 are considered as negative samples. Finally, there are 7,193 negative samples and 628 positive samples in the training dataset of binding affinity predictor.

The expression predictor is also retrained with the samples of Omicron BA.1 and Omicron BA.2 in the DMS dataset from [13]. We normalize the provided expression scores with that of Omicron BA.2 to get the training labels. There are 7,182 negative samples and 639 positive samples in the training dataset of expression predictor.

The exception is that the antibody escape predictor is not retrained since the DMS dataset currently available for antibody escape only involves the wild-type RBD (i.e., the benchmark dataset of antibody escape prediction task).

#### 4.12.2 Intrapandemic evolution prediction

To explore the essential connection between S486P mutation and transmissibility, we mutate the residue on site 486 of XBB.1 into other 19 residues. Subsequently, the binding affinity predictor, expression predictor, and antibody escape predictor are adopted to predict properties of each mutation. We adopt the regression output and conduct normalization within each kinds of property due to the difference of output range between predicted properties for subsequent analysis.

#### 4.12.3 High-risk mutation sites prediction

To recommend high-risk mutation sites of BA.5 and XBB.1.5, we take them as starting sequences, and enumerate all the variants with single-mutation through *in silico* deep mutational scanning. Subsequently, the ACE2 binding affinity, expression and antibody escape of each variant are predicted. As to the antibody used during antibody escape prediction, we adopt BD57-0129 from previous publication [14] as it achieves relatively strong neutralizing capability. For the predicted viral protein properties of variants, we set different thresholds for the probability of the classification output to perform variants screening. Based on previous studies [64–66], partial sacrifice in ACE2 binding affinity is acceptable, while antibody escape is more important for SARS-CoV-2 intrapandemic evolution. Therefore, we set the screening threshold for binding affinity low (0.25 here) and antibody escape moderately (0.5 here). Considering the importance of RBD expression level for the survival of variants, we set the screening threshold for expression higher (0.7 here). After screening the created single-mutation variants with the above thresholds, the number of remaining mutations at each site is considered as indicator of risk level, in which the higher the number of remaining mutations at a site, the higher the risk of this site. Finally, our pipeline recommends the most risky mutation sites based on the number of screened mutations of each site.

In this work, we use AUC to evaluate the ability to predict the potential high-risk mutation sites of different models. Specifically, we consider the wet-lab deep mutational scanning statistic results from [14, 15] as ground-truth (Fig.S3). We set different thresholds from the minimum to the maximum of evolutionary scores, and calculate the true positive rate (TPR) and false positive rate (FPR) with each threshold to draw ROC curve, and AUC is the area under the resulting ROC curve. For our pipeline, we use the number of screened mutations at each site as evolutionary scores. For MLAEP-core pipeline [17], we use the difference of KL-divergence of each site before and after *in silico* evolution as evolutionary scores.

## Data availability

UniRef90 for protein language model pre-training can be downloaded from https://www.uniprot.org/. The raw deep mutational scanning data of binding affinity, expression, and antibody escape prediction tasks are freely available from previous studies [12–14]. The processed data are publicly available at https://github.com/ZhiweiNiepku/E2VD.

## Code availability

Relevant code and models are available at https://github.com/ZhiweiNiepku/E2VD.

## Acknowledgements

We thank Ming Li for his discussion of the protocol and AI for Science (AI4S)-Preferred Program, Peking University Shenzhen Graduate School, China. This work was financially supported by the National Key R&D Program of China (No. 2022ZD0118201), Natural Science Foundation of China (No. 61972217, 32071459, 62176249, 62006133, 62271465, 61825101, 62088102), and R&D Program of Guangzhou Laboratory (Grant No.SRPG22-001).

## Competing interests

The authors declare no competing interests.

## Supplementary information

### S1 Baseline models

Here, we implement several baseline models [17, 19, 22, 24–26] on binding affinity, expression, antibody escape, and high-risk mutation sites prediction tasks. For the baseline model of high-risk mutation sites prediction, we conduct *in silico* evolution using MLAEP [17]. MLAEP adopted genetic algorithm to perform variants population evolution, and the difference of KL-divergence of each site between the original variant population and the evolved variant population can be considered as an indicator of the potential high-risk mutation sites. Therefore, we adopt the difference of KL-divergence before and after *in silico* evolution of each site as evolutionary scores to calculate AUC.

### S2 Protein language model pre-training hyperparameters

**Table S1.**
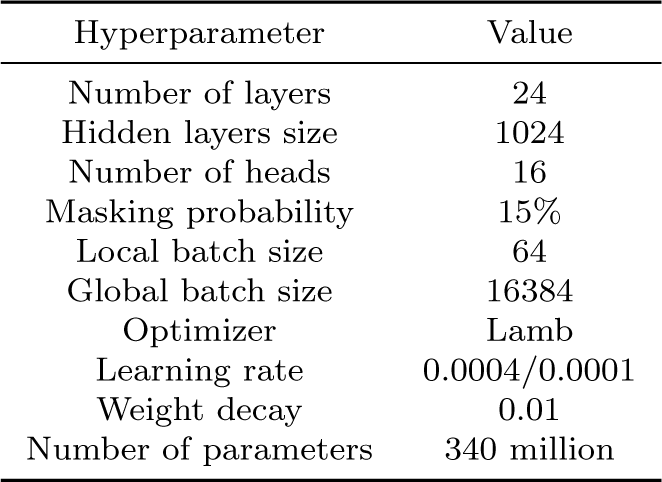
Hyperparameters of customized protein language model.

| Hyperparameter | Value |
| --- | --- |
| Number of layers | 24 |
| Hidden layers size | 1024 |
| Number of heads | 16 |
| Masking probability | 15% |
| Local batch size | 64 |
| Global batch size | 16384 |
| Optimizer | Lamb |
| Learning rate | 0.0004/0.0001 |
| Weight decay | 0.01 |
| Number of parameters | 340 million |

### S3 Ablation experiments of different protein language models

**Table S2.** Results of different protein language models on binding affinity prediction task.

| Model | AUC | Accuracy | F1 score | Precision | Recall | MSE | PCC |
| --- | --- | --- | --- | --- | --- | --- | --- |
| ESM-2(15B) | 0.9383 | 0.9037 | 0.9066 | 0.8769 | 0.9407 | 0.059 | 0.863 |
| ProtT5(3B) | 0.9281 | 0.8741 | 0.8746 | 0.8631 | 0.8889 | 0.079 | 0.818 |
| ESM-1v | 0.939 | 0.8911 | 0.8927 | 0.8806 | 0.9081 | 0.063 | 0.853 |
| E2VD | 0.9298 | 0.9111 | 0.9158 | 0.8743 | 0.963 | 0.058 | 0.87 |

**Table S3.**
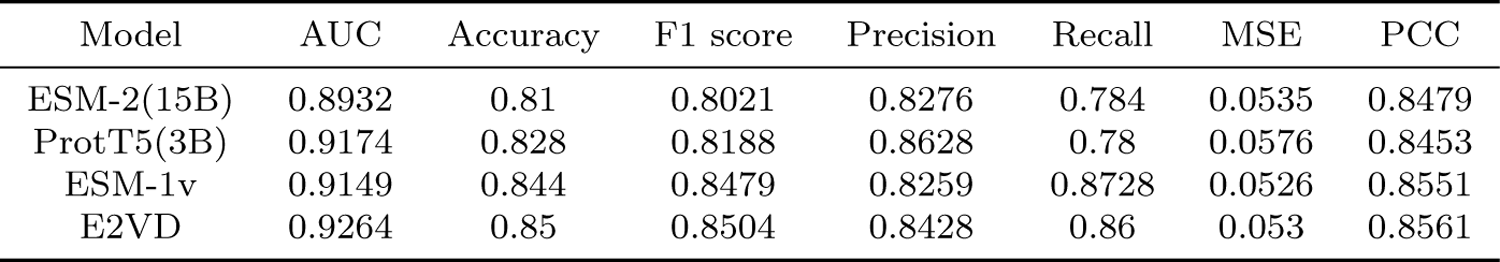
Results of different protein language models on expression prediction task.

**Table S4.** Results of different protein language models on antibody escape prediction task.

| Model | AUC | Accuracy | F1 score | Precision | Recall | MSE | PCC |
| --- | --- | --- | --- | --- | --- | --- | --- |
| ESM-2(650M) | 0.8737 | 0.783 | 0.8061 | 0.7287 | 0.9019 | 0.0425 | 0.6969 |
| ProtT5(3B) | 0.8596 | 0.7785 | 0.7862 | 0.7598 | 0.8146 | 0.0451 | 0.6735 |
| ESM-1v | 0.8641 | 0.7826 | 0.7867 | 0.7726 | 0.8048 | 0.0447 | 0.6798 |
| E2VD | 0.874 | 0.794 | 0.796 | 0.7881 | 0.8042 | 0.0432 | 0.697 |

### S4 Prediction performance of different models

**Table S5.** Results of different models on binding affinity prediction task.

| Model | AUC | Accuracy | F1 score | Precision | Recall | MSE | PCC |
| --- | --- | --- | --- | --- | --- | --- | --- |
| NN_MM-GBSA | - | 0.8041 | - | - | - | 0.2 | 0.79 |
| CNN | 0.6952 | 0.5556 | 0.206 | 0.96 | 0.1185 | 0.1361 | 0.6771 |
| E2VD | 0.9298 | 0.9111 | 0.9158 | 0.8743 | 0.963 | 0.058 | 0.87 |

**Table S6.** Results of different models on expression prediction task.

| Model | AUC | Accuracy | F1 score | Precision | Recall | MSE | PCC |
| --- | --- | --- | --- | --- | --- | --- | --- |
| UniRep_V2 | 0.62 | 0.62 | 0.3866 | 1 | 0.24 | 0.1169 | 0.7546 |
| UniRep_V1 | 0.5496 | 0.50 | 0 | 0 | 0 | 0.2368 | 0.1875 |
| DNAPreceiver | 0.8254 | 0.642 | 0.4492 | 0.6491 | 0.368 | 0.0944 | 0.7209 |
| E2VD | 0.9264 | 0.85 | 0.8504 | 0.8428 | 0.86 | 0.053 | 0.8561 |

**Table S7.**
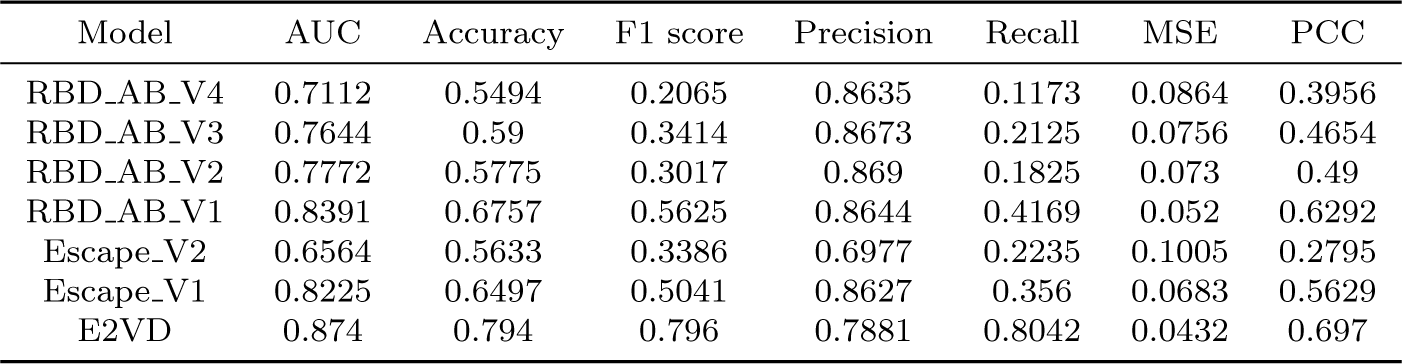
Results of different models on antibody escape prediction task.

| Model | AUC | Accuracy | F1 score | Precision | Recall | MSE | PCC |
| --- | --- | --- | --- | --- | --- | --- | --- |
| RBD_AB_V4 | 0.7112 | 0.5494 | 0.2065 | 0.8635 | 0.1173 | 0.0864 | 0.3956 |
| RBD_AB_V3 | 0.7644 | 0.59 | 0.3414 | 0.8673 | 0.2125 | 0.0756 | 0.4654 |
| RBD_AB_V2 | 0.7772 | 0.5775 | 0.3017 | 0.869 | 0.1825 | 0.073 | 0.49 |
| RBD_AB_V1 | 0.8391 | 0.6757 | 0.5625 | 0.8644 | 0.4169 | 0.052 | 0.6292 |
| Escape_V2 | 0.6564 | 0.5633 | 0.3386 | 0.6977 | 0.2235 | 0.1005 | 0.2795 |
| Escape_V1 | 0.8225 | 0.6497 | 0.5041 | 0.8627 | 0.356 | 0.0683 | 0.5629 |
| E2VD | 0.874 | 0.794 | 0.796 | 0.7881 | 0.8042 | 0.0432 | 0.697 |

### S5 Ablation experiments for standardization

The feature standardization we perform amplifies the differences between the extracted sequence embeddings, which serves a similar purpose to the proposed local-global dependence coupling module. In order to distinguish the impact of feature standardization and the proposed local-global dependence coupling module on prediction performance, we perform ablation experiments on the binding affinity benchmark dataset. As shown in Table S8, without embedding standardization (w/o Std), the model’s prediction performance drops slightly compared to the original complete model (E2VD), with Accuracy and PCC dropping by approximately 5.6% and 9.1%, respectively. When local-global dependence coupling module is not introduced (w/o LG), the prediction performance of the model drops severely compared with the original complete model (E2VD), with Accuracy and PCC dropping by approximately 22.6% and 66.2% respectively. The above results indicate that the local-global dependence coupling module is the main contributor to amplifying the differences of similar sequences, and feature standardization only contributes slightly to it.

**Table S8.** Ablation results for standardization on binding affinity prediction task.

| Model | AUC | Accuracy | F1 score | Precision | Recall | MSE | PCC |
| --- | --- | --- | --- | --- | --- | --- | --- |
| w/o LG&Std | 0.5885 | 0.50 | 0.6667 | 0.50 | 1 | 0.2356 | 0.1302 |
| w/o LG<sup>1</sup> | 0.7007 | 0.6852 | 0.6887 | 0.6832 | 0.6963 | 0.2349 | 0.2077 |
| w/o Std<sup>2</sup> | 0.9248 | 0.8556 | 0.8516 | 0.8734 | 0.837 | 0.093 | 0.7787 |
| E2VD | 0.9298 | 0.9111 | 0.9158 | 0.8743 | 0.963 | 0.058 | 0.87 |

<sup>1</sup>LG represents local-global dependence coupling.
<sup>2</sup>Std represents standardization.

### S6 Convolution kernel size ablation experiments

To choose a suitable kernel size for convolution networks in our model, we train four models with different kernel sizes, including 1, 3, 5 and 7. Overall, the prediction performances on binding affinity prediction task of the four trained models are close under different convolution kernel sizes. Considering that the trained model with a kernel size of 3 achieves the highest classification accuracy, we adopt this size in this work. We don’t conduct further ablation studies on convolution hyperparameters as this is not the key point of this work.

**Table S9.** Ablation results of convolution kernel size.

| Size | AUC | Accuracy | F1 score | Precision | Recall | MSE | PCC |
| --- | --- | --- | --- | --- | --- | --- | --- |
| 1 | 0.932 | 0.90 | 0.9083 | 0.8434 | 0.9852 | 0.0618 | 0.8652 |
| 3 | 0.9298 | 0.9111 | 0.9158 | 0.8743 | 0.963 | 0.058 | 0.87 |
| 5 | 0.938 | 0.90 | 0.9057 | 0.8559 | 0.963 | 0.0551 | 0.8756 |
| 7 | 0.9391 | 0.8926 | 0.8951 | 0.8746 | 0.9185 | 0.0532 | 0.8885 |

### S7 Module ablation experiments

**Table S10.** Module ablation results based on our customized protein language model.

| Model | AUC | Accuracy | F1 score | Precision | Recall | MSE | PCC |
| --- | --- | --- | --- | --- | --- | --- | --- |
| w/o MT&LG | 0.7328 | 0.50 | 0 | 0 | 0 | 0.2219 | 0.385 |
| w/o MT | 0.8406 | 0.5741 | 0.2633 | 0.9714 | 0.1556 | 0.06 | 0.871 |
| w/o LG | 0.7007 | 0.6852 | 0.6887 | 0.6832 | 0.6963 | 0.2349 | 0.2077 |
| Ours(E2VD) | 0.9298 | 0.9111 | 0.9158 | 0.8743 | 0.963 | 0.058 | 0.87 |

**Table S11.**
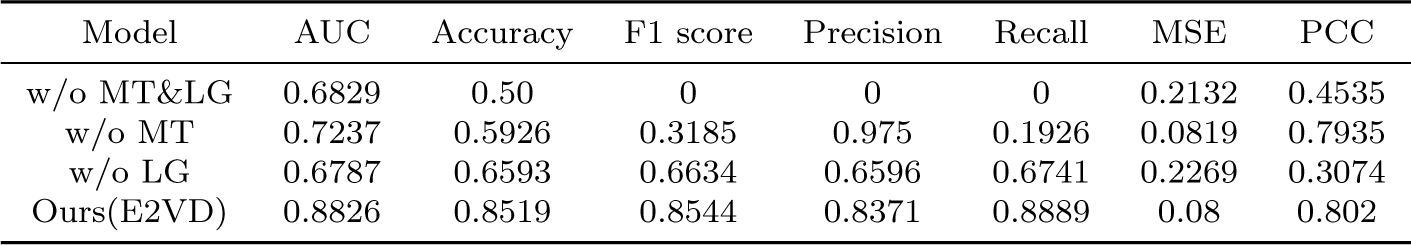
Module ablation results based on ESM-2(650M).

| Model | AUC | Accuracy | F1 score | Precision | Recall | MSE | PCC |
| --- | --- | --- | --- | --- | --- | --- | --- |
| w/o MT&LG | 0.6829 | 0.50 | 0 | 0 | 0 | 0.2132 | 0.4535 |
| w/o MT | 0.7237 | 0.5926 | 0.3185 | 0.975 | 0.1926 | 0.0819 | 0.7935 |
| w/o LG | 0.6787 | 0.6593 | 0.6634 | 0.6596 | 0.6741 | 0.2269 | 0.3074 |
| Ours(E2VD) | 0.8826 | 0.8519 | 0.8544 | 0.8371 | 0.8889 | 0.08 | 0.802 |

**Table S12.**
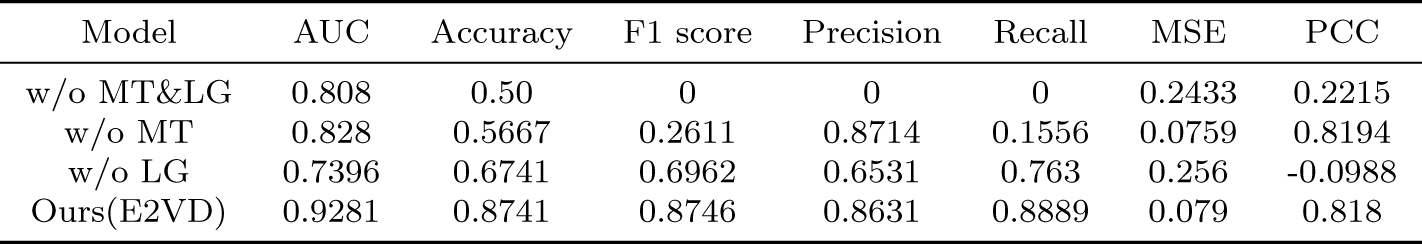
Module ablation results based on ProtT5(3B).

| Model | AUC | Accuracy | F1 score | Precision | Recall | MSE | PCC |
| --- | --- | --- | --- | --- | --- | --- | --- |
| w/o MT&LG | 0.808 | 0.50 | 0 | 0 | 0 | 0.2433 | 0.2215 |
| w/o MT | 0.828 | 0.5667 | 0.2611 | 0.8714 | 0.1556 | 0.0759 | 0.8194 |
| w/o LG | 0.7396 | 0.6741 | 0.6962 | 0.6531 | 0.763 | 0.256 | -0.0988 |
| Ours(E2VD) | 0.9281 | 0.8741 | 0.8746 | 0.8631 | 0.8889 | 0.079 | 0.818 |

### S8 Dimensionality reduction visualization

**Fig. S1.**
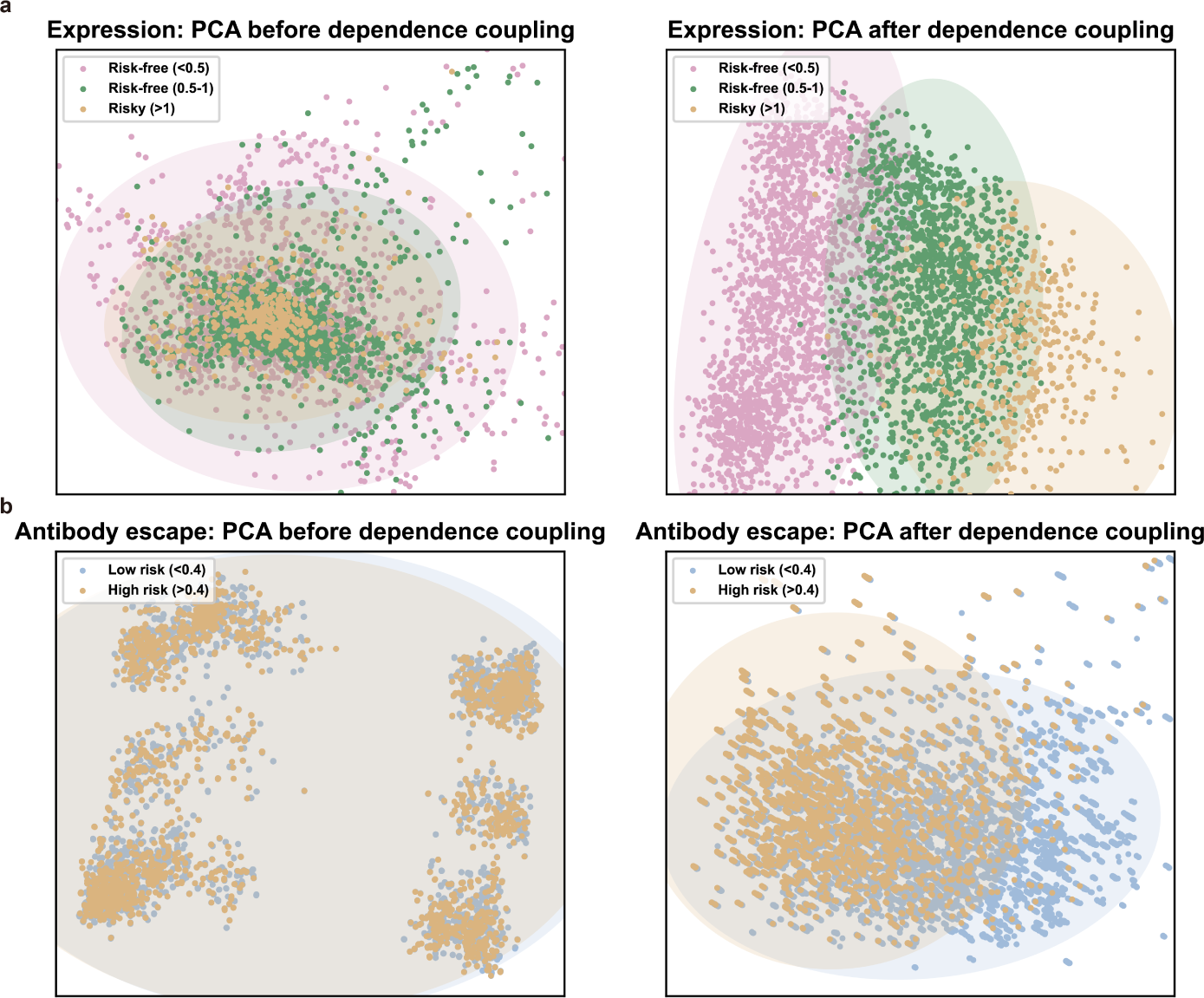
Dimensionality reduction visualization of expression (a) and antibody escape (b) prediction tasks. In expression prediction tasks, three types of mutations are presented, including Risk-free (*MFI ratio<*0.5), Risk-free (0.5*<MFI ratio<*1), and Risky (*MFI ratio>*1). In antibody escape prediction tasks, two types of mutations are presented, including Low-risk (*Escape score<*0.4) and High-risk (*Escape score>*0.4).

### S9 Loss function ablation experiments

**Table S13.**
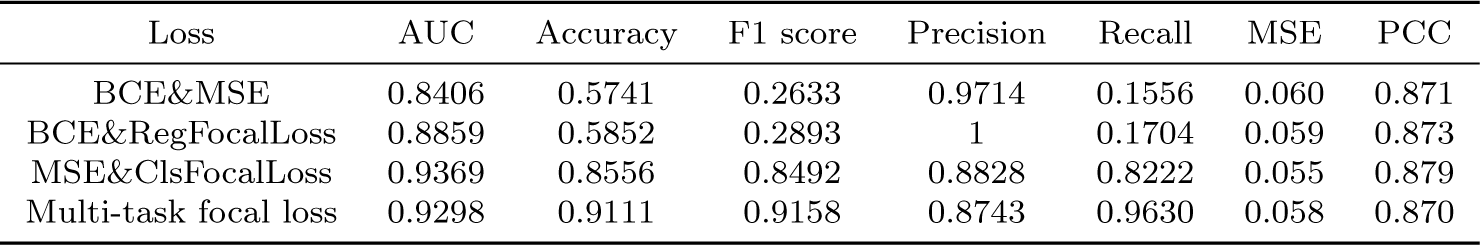
Ablation results of different loss functions.

| Loss | AUC | Accuracy | F1 score | Precision | Recall | MSE | PCC |
| --- | --- | --- | --- | --- | --- | --- | --- |
| BCE&MSE | 0.8406 | 0.5741 | 0.2633 | 0.9714 | 0.1556 | 0.060 | 0.871 |
| BCE&RegFocalLoss | 0.8859 | 0.5852 | 0.2893 | 1 | 0.1704 | 0.059 | 0.873 |
| MSE&ClsFocalLoss | 0.9369 | 0.8556 | 0.8492 | 0.8828 | 0.8222 | 0.055 | 0.879 |
| Multi-task focal loss | 0.9298 | 0.9111 | 0.9158 | 0.8743 | 0.9630 | 0.058 | 0.870 |

### S10 Prediction performance of different loss functions

**Fig. S2.**
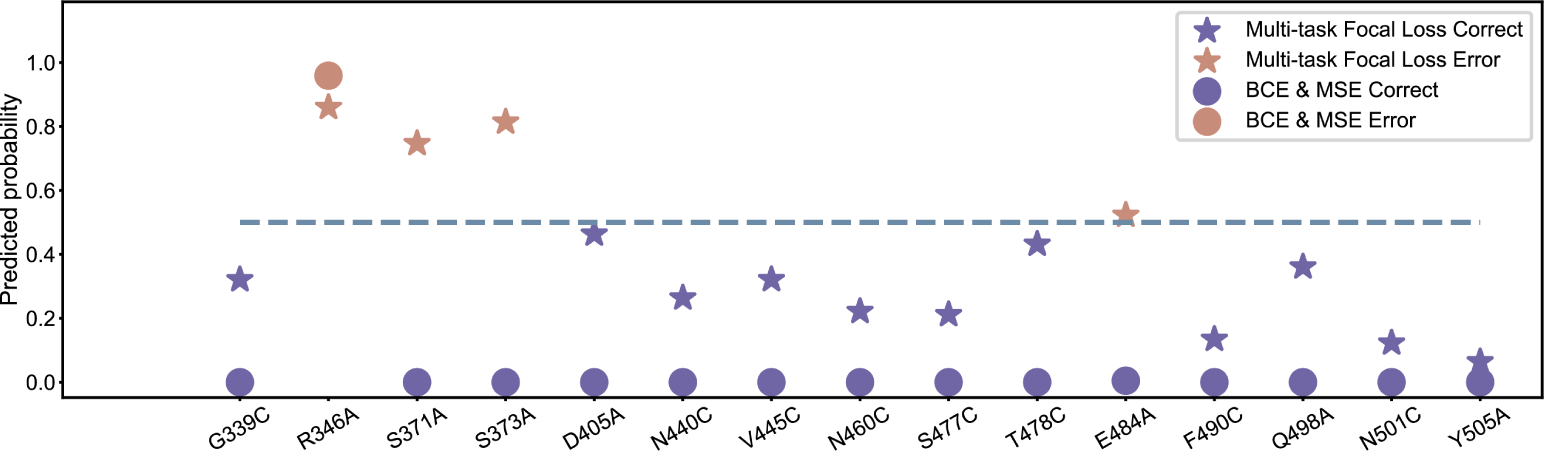
Prediction performance of different loss functions on harmful mutations.

**Table S14.** Overall prediction performance of different loss functions on the comprehensive dataset consisting of both beneficial and harmful mutations.

| Loss | AUC | Accuracy | F1 score | Precision | Recall |
| --- | --- | --- | --- | --- | --- |
| BCE&MSE | 0.7467 | 0.5333 | 0.2222 | 0.6667 | 0.1333 |
| Multi-task focal loss | 0.7867 | 0.7667 | 0.7742 | 0.75 | 0.80 |

### S11 Generalization performance of different models

**Table S15.**
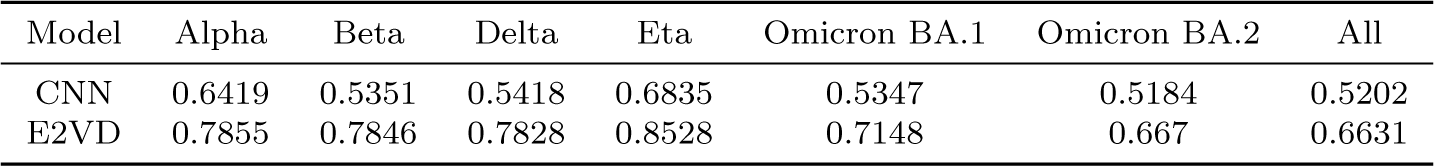
OPPs of six lineages and the mixture of them on binding affinity prediction task.

**Table S16.** OPPs of six lineages and the mixture of them on expression prediction task.

| Model | Alpha | Beta | Delta | Eta | Omicron BA.1 | Omicron BA.2 | All |
| --- | --- | --- | --- | --- | --- | --- | --- |
| DNAPreceiver | 0.7443 | 0.7438 | 0.7139 | 0.7527 | 0.6578 | 0.6125 | 0.5668 |
| UniRep_V2 | 0.535 | 0.5222 | 0.5118 | 0.5317 | 0.5236 | 0.5256 | 0.5163 |
| E2VD | 0.8227 | 0.7886 | 0.7337 | 0.8237 | 0.6743 | 0.5946 | 0.6689 |

### S12 Prediction performance of high-risk mutation sites

**Fig. S3.**
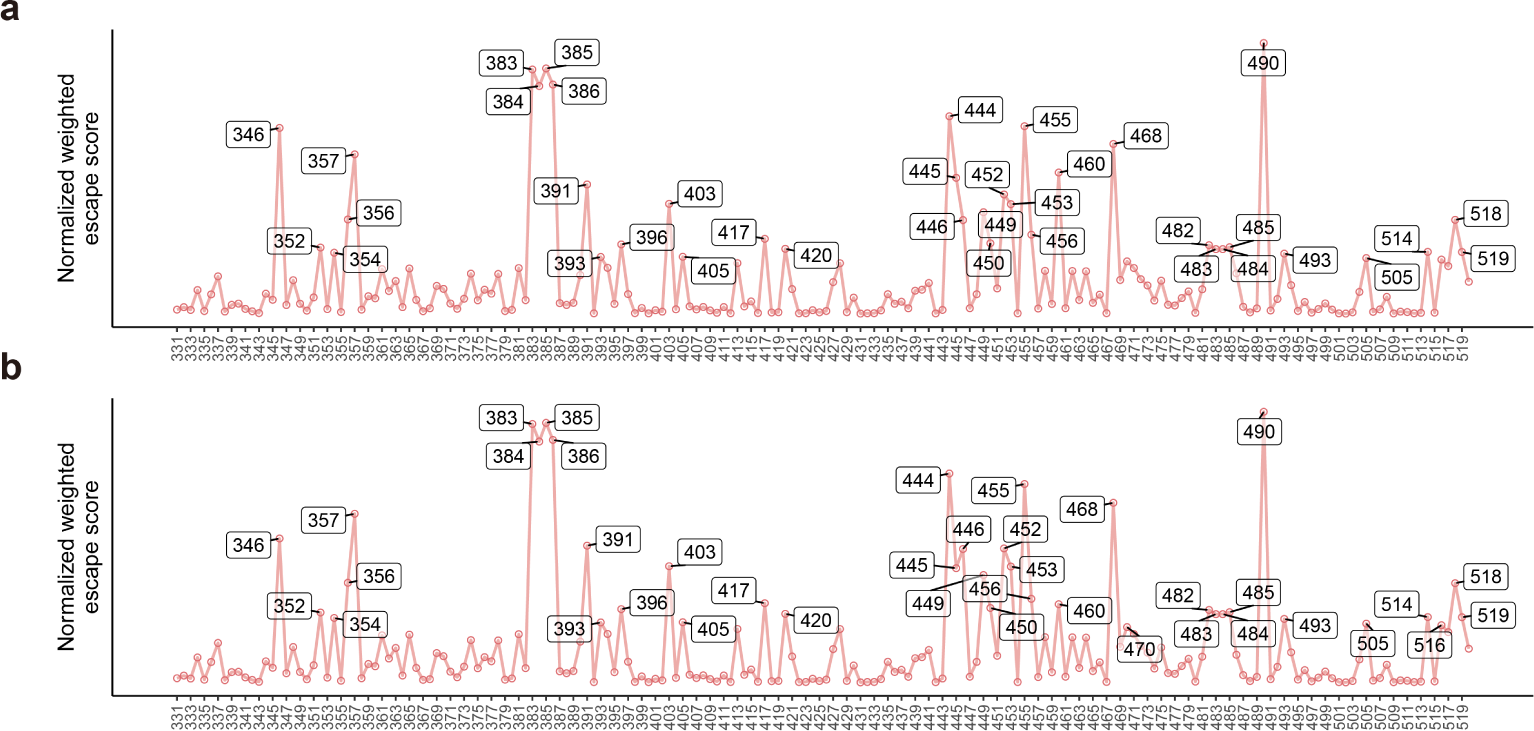
The wet-lab deep mutational scanning statistic results of BA.5 (a) and XBB.1.5 (b). The sites marked in white boxes are the recommended risky ones.

**Table S17.** AUCs of different methods on BA.5 and XBB.1.5.

| Model | BA.5 | XBB.1.5 |
| --- | --- | --- |
| MLAEP | 0.542 | 0.6094 |
| E2VD | 0.724 | 0.661 |

### S13 Model hyperparameters

**Table S18.** Hyperparameters of binding affinity predictor.

| Hyperparameter | Value |
| --- | --- |
| Learning rate | 2e-5 |
| Number of epochs | 150 |
| $\gamma$ | 1 |
| $\beta$ | 0.998 |
| Dropout rate | 0.6 |

**Table S19.** Hyperparameters of expression predictor.

| Hyperparameter | Value |
| --- | --- |
| Learning rate | 2e-5 |
| Number of epochs | 150 |
| $\gamma$ | 1 |
| $\beta$ | 0.99 |
| Dropout rate | 0.6 |

**Table S20.** Hyperparameters of antibody escape predictor.

| Hyperparameter | Value |
| --- | --- |
| Learning rate | 1e-4 |
| Number of epochs | 70 |
| $\gamma$ | 1.2 |
| $\beta$ | 0.88 |
| Dropout rate | 0.6 |

## Footnotes

1 https://outbreak.info/

## Notes

### Competing Interest Statement

The authors have declared no competing interest.

### Summary of Updates

figure revised and supplemental information updated

